# Simulations predict that alterations in balance control contribute to the higher cost of walking with a prosthesis

**DOI:** 10.64898/2026.09.03.749211

**Authors:** Wouter Muijres, Maarten Afschrift, Renaud Ronsse, Tom Van Wouwe, Friedl De Groote

## Abstract

The high metabolic cost of walking in individuals with a transtibial amputation is often attributed to reduced ankle propulsion. However, the cost of stabilizing walking may also increase after amputation, potentially explaining why restoring ankle propulsion does not always normalize metabolic cost. Because the contributions of propulsion and stabilization are difficult to dissociate experimentally, we used simulations to probe the effects of transtibial amputation and different prostheses on walking cost. We simulated gait using a conceptual planar torque-driven model. Leg joint torques were controlled by feedforward and linear, time-varying full state feedback. We computed feedforward torques associated with propulsion and feedback gains for stabilizing walking that minimized expected effort (sum of torques squared) in the presence of sensorimotor noise while imposing stability. We simulated walking for an intact model and three walkers with a prosthesis: a passive, a feedforward-controlled active, and a feedforward and local feedback-controlled active prosthesis. We solved a deterministic approximation of the resultant stochastic optimal control problems for different levels of sensorimotor noise. Feedforward and expected feedback torques were higher for the passive prosthesis walker than for the intact walker. An active prosthesis restored total biological effort (i.e., effort of all but the prosthesis joint) to the level of the intact walker, but this effort was generated by fewer biological joints. Biological feedback effort was higher in prosthesis walkers than in the intact walker. Adding local feedback to an active prosthesis reduced biological feedback effort, but only at low noise levels. Our results suggest that walking with passive prosthesis increases effort related to both propulsion and stabilization. While active prostheses can support propulsion, local feedback control only marginally reduced stabilization-related effort, suggesting that controllers require sensory information from the rest of the body to fully restore the metabolic cost of walking.

**Author summary:** Amputation of the lower leg substantially increases the energy cost of walking, reducing mobility. Loss of ankle propulsion is often considered the main cause, motivating the development of active prostheses that generate propulsion. However, restoring propulsion does not consistently reduce the energy cost of walking, suggesting other contributing factors. Stabilizing walking also requires energy, and recent studies suggest that the required energy may be increased after amputation. Because an amputation simultaneously affects propulsion and balance, experimentally disentangling contributions of reduced ankle propulsion and altered balance control to increased energetic cost is hard. We therefore used simulations to study how sensorimotor deficits due to amputation affect energy costs of propulsion and balance control. Our simulations suggest that the high energy cost of walking with a passive prosthesis results from increased effort for both propulsion and balance. While an active prosthesis can restore effort required for propulsion, it does not restore effort required for balance control, even when supporting balance with local feedback of prosthetic ankle kinematics. These findings suggest that the limited ability of existing active prostheses to reduce walking energy cost may - at least partially - be due to the inability to reduce the energy cost for balance control.

## Introduction

Individuals with a transtibial amputation consume, on average, 12% (non-vascular) to 36% (vascular) more metabolic energy than individuals without an amputation (1). This higher metabolic cost is thought to arise from a loss of ankle power (2,3). However, active prostheses that restore ankle power do not necessarily decrease metabolic cost (4), suggesting that there are other reasons for the observed increase in metabolic cost. Transtibial amputation also affects the ability to stabilize walking, resulting in a higher fall incidence in individuals with a transtibial amputation (5). Walking is stabilized by continuous sensorimotor feedback (6), but after amputation, both sensory information from and direct control of the missing limb are lacking, leading to alterations in stabilizing control strategies (7–11). As there is a metabolic cost associated with stabilizing walking (12–14), alterations in control strategies might influence the metabolic cost of stabilizing walking. It is hard to experimentally dissociate between increases in metabolic cost due to a loss of ankle power and due to altered balance control, as both occur simultaneously and interact. Therefore, we used predictive simulations of locomotion to probe the effect of sensorimotor deficits due to amputation on the contribution of alterations in propulsion and stabilization to the cost of walking with a prosthesis.

Transtibial amputation leads to an increase in walking metabolic cost (1) that cannot be fully explained by a lack of ankle power. Limiting ankle push-off power increases the metabolic cost of walking in individuals without an amputation (15). Analyses based on inverted pendulum models of walking have shown that ankle push-off can reduce energy dissipation during collision (16). This has been confirmed by experimental observations. A higher metabolic cost of walking has been associated to a higher dissipation of mechanical energy during step-to-step transition in individuals with and without an amputation (2,17), and individuals with a transtibial amputation dissipate more energy during step-to-step transition than individuals without an amputation when walking (2). However, differences in energy dissipation during step-to-step transitions only partially explain differences in metabolic cost of walking between individuals with and without an amputation (2), suggesting that the increase in metabolic energy is not only due to reduced ankle power. This is in line with the variable effect of active ankle prostheses that aid propulsion by restoring ankle power on the metabolic cost of walking. Some studies report that such active prostheses can reduce the metabolic cost of walking (3,18). Yet, other studies observed that active ankle prostheses fail to restore metabolic energy consumption (4,19,20), even when the prosthesis generates large push-off powers with optimized timing (4). A lack of ankle power may thus not be the only contributor to the higher metabolic cost of walking in individuals with a transtibial amputation.

There is a metabolic cost associated with stabilizing walking, and this cost might be higher in individuals with an amputation. In individuals without an amputation, external stabilization of the pelvis decreases the metabolic cost of walking (12), whereas visual (13,21) and mechanical perturbations (22) increase the metabolic cost of walking. In individuals with a dysvascular transtibial amputation, handrail support decreases the metabolic cost of walking on a treadmill (23). However, it is unclear whether the metabolic cost of stabilizing walking differs between individuals with and without an amputation. Ijmker et al. (24) found no difference in the decrease in metabolic cost upon frontal plane stabilization of the pelvis in individuals with and without a transtibial amputation. In contrast, we recently found that individuals with a transtibial amputation (mostly traumatic etiology) consume more energy than individuals without an amputation to stabilize walking against sagittal plane treadmill speed perturbations (10). Whereas individuals without a transtibial amputation modulate their ankle torque based on sensorimotor feedback in response to sagittal plane perturbations (6), individuals with a transtibial amputation cannot use this strategy on the amputated side. This might explain why they rely more on step length adjustments when perturbed while standing on their prosthesis (10).

Simulations can help dissociate the effects of a lack of ankle power and altered sensorimotor control on the metabolic cost of walking, but existing simulations of walking with a prosthesis do not capture how walking is stabilized against uncertainty. Many previous simulations, which either tracked experimental kinematics (often of individuals without an amputation) (25–27) or predicted the kinematics without relying on experimental data (28–30), yielded similar metabolic costs for walking with a passive prosthesis as for walking with biological limbs. Hence, these simulations do not capture the higher energetic cost of walking that is observed in most individuals with a transtibial amputation walking with a passive prosthesis (1). In addition, simulations based on a planar model (seven degrees of freedom, 16 muscles) predicted that powered prostheses can reduce the metabolic cost of walking to a level below that of individuals without an amputation (29,30), whereas observed metabolic benefits due to active prosthesis use have been variable and modest at best (4,19,31). These simulations predicted highly asymmetric gait patterns, whereas increased symmetry in terms of net leg work has been observed when walking with an active versus passive prosthesis (3). A shortcoming of these prior simulations is that they did not account for uncertainty, e.g., due to sensorimotor noise, and therefore also not for the metabolic cost of stabilizing walking against uncertainty. Instead, they solved for muscle inputs that minimized a movement-related cost (e.g., metabolic energy) and sometimes deviations from a desired walking pattern, without modeling the underlying control policy.

Simulations of movement that account for uncertainty capture important features of motor control but are computationally challenging. Whereas uncertainty is typically not accounted for when simulating walking, it has been more common to account for uncertainty in simulations aiming at understanding motor control of reaching (32–35). The resulting optimal control policies lead to simulations that capture important features of human behavior, such as smoothness of movement trajectories (32), the minimal intervention principle (i.e., the accumulation of variability in task-irrelevant dimensions) (33,35), and the trade-off between movement speed and accuracy (32). Most prior optimal control simulations that accounted for uncertainty were based on very simple and often linear models of musculoskeletal dynamics for computational reasons (32,34,36). However, walking dynamics is highly non-linear, which might explain why simulations of walking that account for uncertainty are rare. Koelewijn and van den Bogert (37) simulated walking in the presence of noise using a planar 9-degree-of-freedom torque-driven model. However, their control policy was overly simple, possibly for numerical reasons. Their model was controlled by joint level proportional-derivative feedback whereas human balance control relies on task level feedback, e.g. ankle torque modulations can be explained by feedback from center of mass kinematics but not by feedback from local joint angles and velocities (6). We recently developed a computationally efficient approach to simulate non-linear dynamics in the presence of noise (35). By approximating the resulting stochastic optimal control problem by a deterministic problem (35), we could leverage recent advances in computational methods (e.g., algorithmic differentiation and direct collocation) to simulate movement in the presence of sensorimotor noise. By applying this approach to standing balance (35), we demonstrated that sensory deficits lead to alterations in the minimal effort sensorimotor feedback control strategy to stabilize standing.

Here, we use our approximate stochastic optimal control framework to simulate intact walking and walking with different prostheses in the presence of sensorimotor noise. We evaluated the effect of walking with a prosthesis on the cost of propulsion and stabilization of walking by comparing simulations of walking without an amputation and walking with a passive prosthesis, a feedforward-controlled active prosthesis, and an active prosthesis with feedforward and local feedback control. All simulations were based on conceptual torque-driven five-segment models. Torques were controlled by feedforward and full-state linear feedback control. Feedforward torques in the absence of noise generate the mean walking pattern and therefore mainly reflect propulsion, whereas feedback torques stabilize walking based on deviations from the mean walking pattern. We hypothesized (i) that walking with a passive prosthesis would result in higher feedforward torques and higher feedback torques, (ii) that active prostheses would be successful at reducing the feedforward torques, but (iii) that only an active prosthesis with local feedback would also reduce feedback torques. Given the use of a conceptual model, we expected simulations to qualitatively but not quantitatively capture human behavior.

## Methods

We simulated prosthetic and intact walking by performing stochastic optimal control simulations based on a planar torque-driven skeletal model.

### Skeletal model

All simulations were based on the same skeletal model with five segments: a trunk, two upper legs, and two lower legs (Figure 1A). Segment lengths and inertial parameters were chosen to mimic the proportions of a human adult (Table 1). At any instant in time, only one leg - the stance leg - was in contact with the ground, whereas the other leg - the swing leg - moved freely above the ground. The stance leg was assumed to be fixed to the ground through a hinge representing the ankle joint. The model thus had five degrees of freedom: the stance ankle angle, two knee angles, and two hip angles. The state (*x*) of the model was described by five joint angles (*θ*) and angular velocities (*ω*). When the endpoint of the swing leg struck the ground, the swing leg became the stance leg and vice versa. The collision was impulsive and preserved angular momentum, i.e., the angular momentum around the former stance ankle equals the angular momentum around the new stance ankle. The formulation of the skeletal dynamics was inspired by (38). We assumed that prosthetic use did not affect the inertial properties of the shank and that the prosthesis was rigidly attached to the shank, and therefore, skeletal dynamics did not differ between the intact and prosthesis walkers.

**Figure 1.**
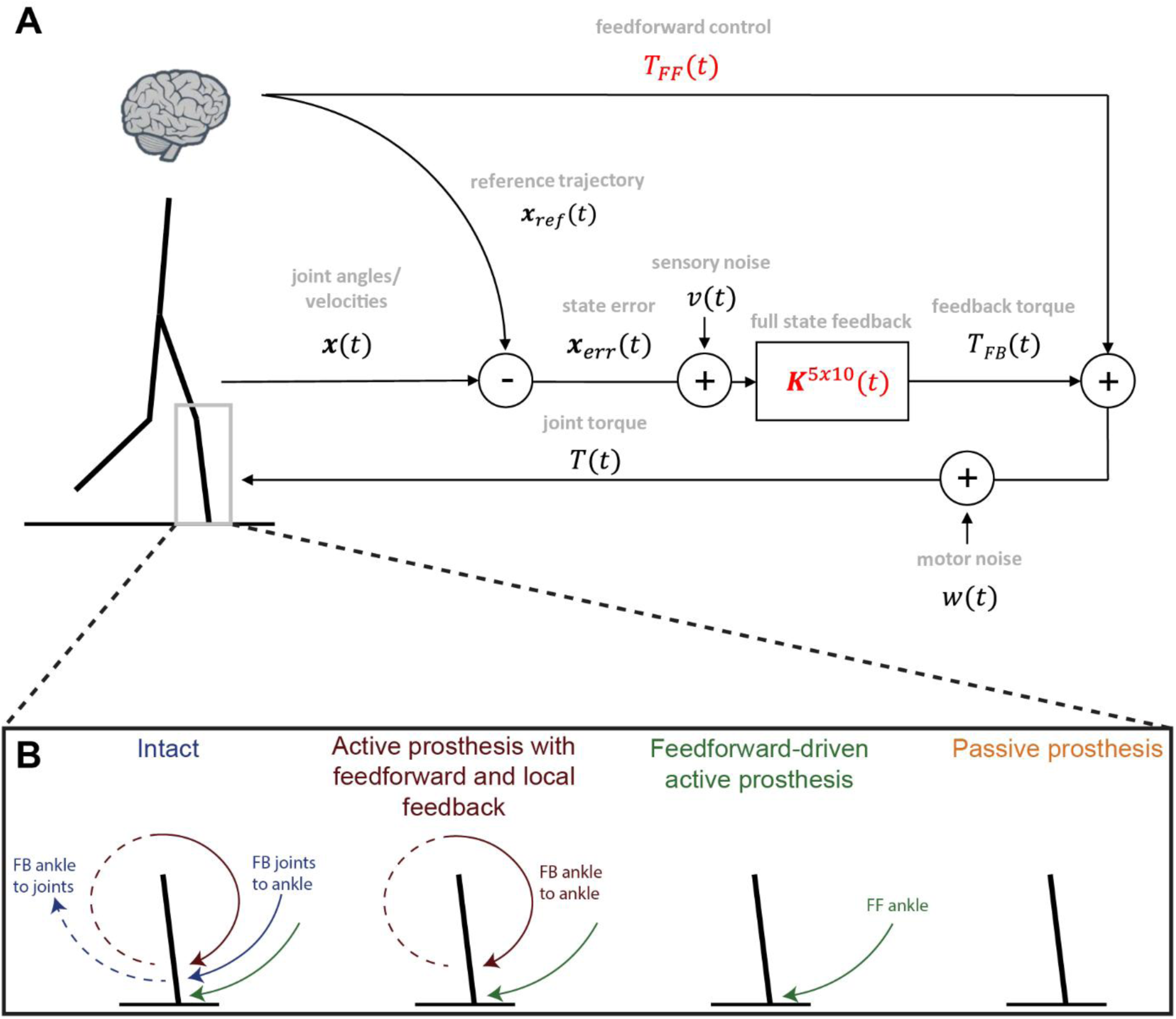
Neuromusculoskeletal model of gait. A. The skeletal system consists of five segments. Joints are driven by torques. At any instance, one single leg is in contact with the ground (i.e., no double support phase), and foot-ground contact was modeled as an impulsive collision. Joint torques consist of a feedforward torque, T_FF_, and a feedback torque, T_FB_. Feedback torques are modeled by linear time-varying full state feedback. Position and velocity errors are computed with respect to the reference state x_ref_, x_err_, and corrupted by additive zero-mean Gaussian noise, v.The reference state trajectory is the state trajectory that results from applying the feedforward torques in the absence of noise and is thus fully determined by the feedforward torques and the system dynamics. The matrix K describes the feedback gains. The total torque, T, is corrupted by additive zero-mean Gaussian noise, w. Red variables, T_FF_ and K, were obtained by solving the optimal control problem. The model was adapted from (_38_). B. We modeled intact walking and walking with three different prostheses (from left to right): an active prosthesis with feedforward and local (ankle angle and angular velocity) feedback control, a feedforward-driven active prosthesis, and a passive prosthesis (i.e., a pin joint for this simple model).

**Table 1.** Inertial properties of the skeletal model. The model was symmetrical. The center of mass of the trunk, upper leg, and lower leg were defined as the distance from the hip, knee, and ankle along the trunk, upper leg, and lower leg segment, respectively (inspired by a scaled OpenSim gait 2392 model (39,40)).

| Segment | mass [kg] | length [m] | center of mass [m] | inertia [kg m<sup>2</sup>] |
| --- | --- | --- | --- | --- |
| trunk | 46 | 0.625 | 0.365 | 2.93 |
| upper leg | 9.3 | 0.40 | 0.22 | 0.14 |
| lower leg | 5 | 0.43 | 0.20 | 0.1 |

### Control policy

In the intact model, a torque actuator drove each degree of freedom. Torques were controlled by both feedforward and feedback contributions and were corrupted by zero-mean Gaussian motor noise *w*(*t*):

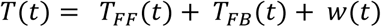

with *T*(*t*), *T_FF_* (*t*), and *T_FB_*(*t*) 5×1 vectors of respectively total torques, feedforward torques, and feedback torques (Figure 1A). We modeled the feedback torque as linear, time-varying feedback of the error between the state trajectory and a reference state trajectory. This error was corrupted by zero-mean Gaussian sensory noise *v*(*t*):

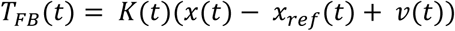

with *K*(*t*) a 5×10 matrix of feedback gains and *x*(*t*) the 10×1 state vector. The reference state trajectory *x_ref_*(*t*) is the state trajectory that results from applying the feedforward torques in the absence of noise. The reference trajectory is thus fully determined by the feedforward torques and the system dynamics.

We modified the actuation and controller of the intact model to represent three types of prostheses (Figure 1B). To model the use of a passive prosthesis, we removed the torque driving and the sensory information from one ankle (i.e., by setting the corresponding gains in K to zero). This simplification does not capture that passive prostheses typically have ankle stiffness and damping, but this could not be implemented in our simple model because it does not explicitly model the feet, and therefore the angle between the shank and foot. To model the use of an active prosthesis without feedback control, we removed the feedback torque for one ankle as well as the contribution from sensory information from this ankle to the feedback torques driving other joints, again by setting the corresponding gains in K to zero. In other words, the prosthesis was controlled by a feedforward torque only. In addition, we tested an active prosthesis with both feedforward and local (ankle angle and angular velocity) feedback control. To this end, we removed feedback from all joints but the ipsilateral ankle to the ankle on one side and feedback from that ankle to all other joints from the intact model. We chose this scenario because the prosthetic ankle’s position and velocity can easily be obtained from the prosthesis encoders, and such local feedback has been used previously in real-world devices (41,42). We were interested in whether such a PD controller could help stabilize walking against sensorimotor noise. We assumed that the feedforward prosthetic torque was noise-free with noisy information from encoders. We assumed the same noise level for prosthetic and biological joint angles and angular velocities.

### Optimal control

We solved for control parameters, i.e., feedforward torque trajectories (*T_FF_* (*t*)) and time-varying feedback gains (*K*(*t*)), that minimized the expected effort while imposing a mean step time of 0.8 s and a mean step length of 0.5 m corresponding to a mean speed of 0.6 m/s, periodicity over a stride, and foot clearance. Because the conceptual model had no feet, foot clearance was imposed by constraining the swing ankle to be above the ground (vertical position larger than 0) from 10% to 90% of the stance time. In humans, muscles consume metabolic energy to generate the mechanical power that is required for walking. The joints in the conceptual model were not actuated by muscles but by ideal torque actuators. Therefore, we used the sum of the squared total (feedforward and feedback) torques as a measure of effort instead of metabolic energy. Active ankle prosthesis effort was included in the optimization of the expected effort under the assumption that the control capacity of active prostheses is limited by a finite power source (e.g., a wearable battery pack). Given that all models had similar skeletal dynamics, differences in expected effort between models are due to differences in the control policies (combination of feedforward torques and feedback gains).

To test the effect of noise on the control policy and resulting movement, we solved for optimal controllers for different levels of sensory noise (see Table 2) with constant motor noise (with a variance of 1^2^ (Nm)^2^s). Note that this continuous-time variance was divided by the length of the integration interval (in seconds) when performing numerical integration (35). The level of motor noise was inspired by previous studies that estimated torque variability from experimental torques during a torque matching task (43,44). There is no experimental data for the uncertainty on angular positions and velocities during walking, and, therefore, we performed simulations at different sensory noise levels. The range of sensory noise levels was chosen based on previous simulations of standing balance control using the stochastic optimal control framework, which resulted in postural sway levels in agreement with experimental observations (35). Note that our model makes an abstraction of the physiological processes underlying state estimation from different sensory inputs (e.g., proprioceptors embedded in the muscles, visual and vestibular information) and that sensory noise should be interpreted as the uncertainty on the joint positions and velocities after state estimation (e.g., in the cerebellum (45)). We assumed that noise was independent for the different joints. In addition, we assumed that position and velocity noise were independent. Hence, all noise covariance matrices were diagonal.

**Table 2.** Joint angular position and velocity sensory noise parameters. For each model, we performed five optimal control simulations to find control policies that minimized expected effort for different levels of joint angular position (θ_joint_) and velocity (θ̇_joint_) noise with a variance of σ^2^

| Noise levels | 1 | 2 | 3 | 4 | 5 |
| --- | --- | --- | --- | --- | --- |
| $\sigma_{\theta_{joint}}^2 ((^\circ)^2 s)$ | $0.01^2$ | $0.02^2$ | $0.05^2$ | $0.1^2$ | $0.2^2$ |
| $\sigma_{\dot{\theta}_{joint}}^2 \left( \left( \frac{^\circ}{s} \right)^2 s \right)$ | $0.01^2$ | $0.02^2$ | $0.05^2$ | $0.1^2$ | $0.2^2$ |

### Solving the optimal control problems

The system dynamics were stochastic due to the presence of motor and sensory noise. We solved the resulting stochastic optimal control problems using our recently developed framework (35). Our framework relied on a deterministic approximation of the stochastic optimal control problems. We approximated the state distribution by a Gaussian distribution. As a result, the state distribution can be described by the mean state (*x̅*) and the covariance matrix (*P*). The dynamics of the mean state were described by setting the noise to zero in the dynamic equations. Note that the reference state equals the mean state. The dynamics of the state covariance were described by the continuous Lyapunov differential equations based on a local first-order approximation of the nonlinear system dynamics around the mean state. To also capture the variability in contact timing, we used the saltation matrix to propagate the covariance matrix at the time of contact (46). The intact walker was symmetric, and we therefore solved for a step while imposing left-right symmetry by constraining the mean state and state covariance of the stance and swing leg after impulsive contact to be equal to the initial mean state and state covariance of the swing and stance leg, respectively. As the prosthetic walkers were asymmetric, we solved for a full stride of the prosthetic walkers and imposed periodicity by constraining the mean state and state covariance after heel strike of the initial stance leg to the initial mean state and state covariance. We approximated stochastic constraint functions by a Gaussian distribution and imposed that the chance of fulfilling the constraint should be 99.7%, e.g., foot clearance was obtained for 99.7% of the trajectories at each time instant. Note that it is impossible to impose that the constraints should always be fulfilled due to the infinitely long tails of a Gaussian distribution.

We solved the approximate deterministic optimal control problems using direct collocation with implicit formulations of the dynamics of the mean state, the dynamics of the state covariance, and the integration scheme. We used a trapezoidal integration scheme with mesh intervals of 0.01 s. We used CasADi to perform automatic differentiation and solved the resulting sparse nonlinear programming problems with IPOPT (47,48).

### Outcome variables

We evaluated the effect of walking with a prosthesis on the expected effort by comparing expected effort over a stride (i.e., time integral of the sum of expected torques squared) of the optimal solutions between the prosthesis and intact walkers. We used feedforward effort (sum of feedforward torques squared) as a measure of effort related to propulsion and feedback effort (sum of expected feedback torques squared) as a measure of effort related to stabilizing walking. Expected effort from the biological joints (i.e., excluding the active prosthesis) was summed to obtain an estimate of total biological effort. Prosthetic walkers had five biological joints (i.e., the hips, knees, and the intact ankle), while the intact walker had six biological joints (i.e., the hips, knees, and the two ankles). To enable a comparison between the common biological joints of intact and prosthetic walkers, we also evaluated the effort of a biological ankle joint and of the other joints separately for the intact walker.

In addition to effort outcomes, we evaluated differences in control strategies (feedback gains) and walking kinematics (mean and standard deviation) between intact and prosthetic walkers. The feedback gains in our models varied over time, e.g., 50 time-varying gains (5 joint torques x 10 states) for the intact walker and 32 time-varying gains for the passive prosthesis walker (4 joint torques x sensory input from 8 states). Therefore, to simplify the comparison between models, we computed root mean square values of the gains.

## Results

Solving the optimal control problems yielded a control strategy consisting of feedforward torques and feedback gains, as well as the corresponding expected effort and kinematic trajectories. Below, we describe observations based on comparing optimal solutions for the intact walker and different prosthesis walkers. Figures that present kinematics and feedback gains show data for the median sensory noise level of 0.05^2^(°)^2^s and 0.05^2^ (°/s)^2^s, as we found that the agreement between simulated data and experimental observations (i.e., variability in fore-aft center of mass position and ankle joint torques) was similar for different sensory noise levels (see Supplemental material S1) but we provided results for the other noise levels in the supplement (Supplemental material S4).

The expected biological effort was 7% to 13% (lowest to highest sensory noise level) higher for the walker with a passive prosthesis than for the intact walker (Figure 2A). The expected biological effort of active prosthesis walkers was more similar to that of the intact walker. Compared to the intact walker, the expected biological effort in the feedforward-controlled active prosthesis walker was similar at low and 5% higher at high sensory noise levels. Expected biological effort in the active prosthesis walker with feedforward and local feedback control was 3% lower at low and 3% higher at high sensory noise levels than the expected biological effort in the intact walker. At low sensory noise levels, local feedback control thus reduced the expected effort compared to the intact and the active prosthesis walker with solely feedforward control. Yet, these differences in expected biological effort between the intact and active prosthesis walkers were small. Therefore, our simulations suggest that walking with an active prosthesis, even with local feedback, requires a similar overall biological effort as walking with two intact limbs. As the intact walker has six biological joints whereas the prosthesis walkers have only five biological joints, this implies that the remaining biological joints (hips, knees, and intact ankle) have to generate larger torques when walking with an active prosthesis than when walking with two intact limbs (Figure 2A, green, yellow, red traces versus light blue trace). The torque generated by the active prostheses, which is free from a biological perspective, can thus not remove the need for compensation by the remaining biological joints after amputation.

**Figure 2.**
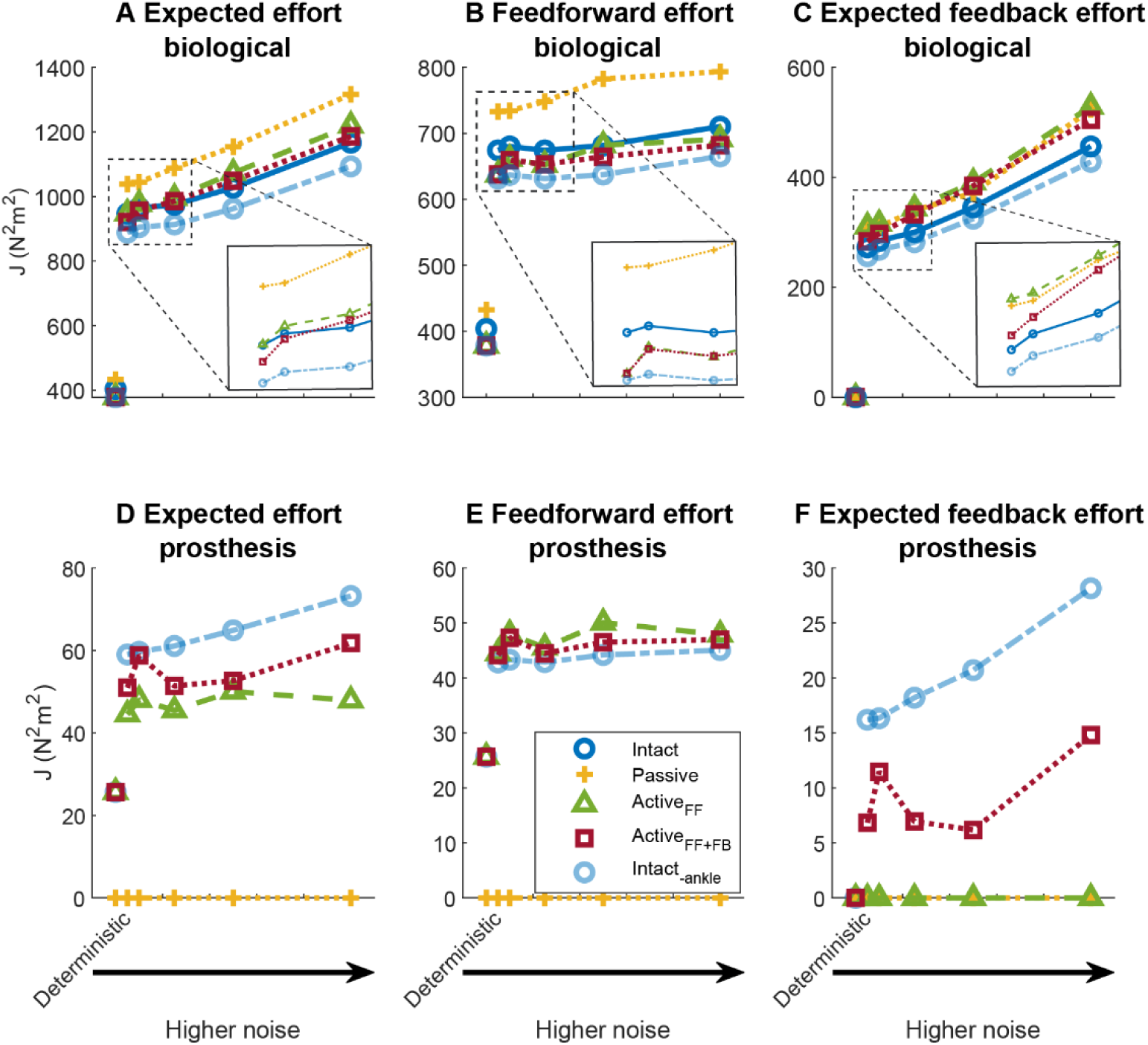
Expected effort from biological joints (A-C) and expected prosthesis effort (D-F) over a stride, when walking with different levels of sensory noise. (A) Expected effort for the intact (blue) and passive prosthesis (yellow) walkers, and for a walker with an active prosthesis that is solely feedforward controlled (green) or feedforward and local feedback controlled (red). Intact-ankle presents (expected) effort of the intact walker minus the (expected) effort of one of the ankles. Therefore, the biological effort of the same five joints is included in Intact_-ankle_ and in the prosthesis walkers. Expected effort is the sum of (B) feedforward effort (feedforward torques squared) and (C) expected feedback effort (expected feedback torques squared). The inserted smaller plots in Panels A-C enlarge the results for the different walkers for the first three sensory noise levels to better visualize differences between walkers at these low noise levels. Panels D-F show the prosthesis effort. Intact_-ankle_ refers to the effort from the biological ankle joint in the intact walker that was controlled based on feedforward and time-varying full-state linear feedback. This was the effort that was not accounted for in the calculation of effort in A-C for Intact_-ankle_. Deterministic on the x-axis denotes the effort for an optimal control solution without sensorimotor noise.

The expected biological feedback effort was higher in all prosthesis walkers than in the intact walker. Walking with a passive prosthesis increased feedforward effort by 7% to 15% and feedback effort by 8% to 15%, depending on the sensory noise level (Figure 2B-C). Simulated increases in expected biological effort when walking with a passive prosthesis were thus due to both increases in feedforward effort and expected feedback effort. Compared to the intact walker, biological feedforward effort in the feedforward-controlled active prosthesis walker was 0% to 6% lower (Figure 2B), and the expected feedback effort was 11% to 16% higher (Figure 2C). A feedforward-controlled active prosthesis may thus reduce feedforward but not feedback effort. Also, the feedforward and local feedback-controlled active prosthesis reduced feedforward effort compared to the intact walker (3% to 6%). While the expected feedback effort in the feedforward and local feedback-controlled active prosthesis was still higher than for the intact walker (4% to 11%), the expected feedback effort at low sensorimotor noise levels was not as high as for the other prosthesis walkers (Figure 2C). This suggests that local feedback control in an active prosthesis normalized expected feedback effort at low but not at high sensory noise levels. In addition, we observed that the active prosthesis generated slightly more feedforward effort than the biological ankle (Figure 2E), whereas the biological ankle generated more feedback effort than the active prostheses (Figure 2F).

Simulated joint angle trajectories were similar for all walkers, but joint angles were more variable for the passive prosthesis walker across sensory noise levels and for all prosthesis walkers at high sensory noise levels (Figure 3). Knee joint angles on the intact and prosthesis side were more variable in the passive prosthesis walker than in the intact walker and active prosthesis walkers. Differences in kinematics between the intact walker and the prosthesis walkers increased with sensory noise levels (Figure S5 and Figure S6). At the highest noise level, the intact walker flexed the knee more over the swing phase than the prosthesis walkers and swing hip and knee angle variability was larger for the prosthesis than for the intact walkers (Supplementary material S4). Differences in joint angular velocities – mean and variability – between walkers followed the same trends as differences in joint angles (Supplementary material S2).

**Figure 3.**
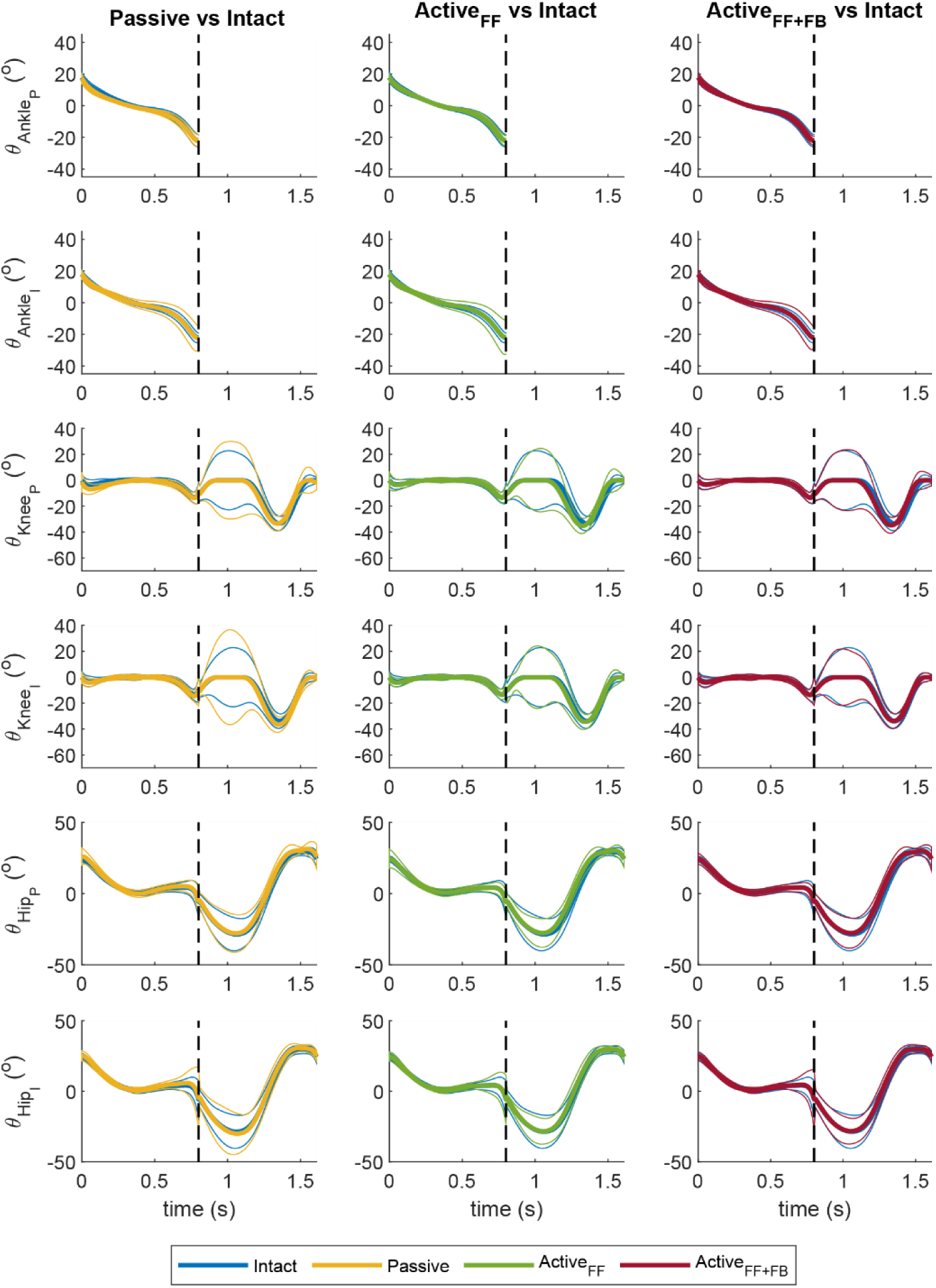
Joint angular positions (θ) over a stride for the prosthesis walkers versus the intact walker at a sensory noise level of 0.05^2^ (°)^2^s and (°/s)^2^s. The optimal control solutions of the passive prosthesis are shown in yellow, of the feedforward controlled active prosthesis in green, the feedforward and feedback-controlled active prosthesis in red, and the intact walker in blue for the intact (I) and the prosthetic (P) legs. Joint angular positions are displayed for the intact and prosthetic ankles, knees, and hips. Thick and thin lines represent the mean and the mean +/- 1 standard deviation, respectively. The dashed vertical line indicates the moment of toe-off, separating the stance phase from the swing phase.

Feedback gains were higher for the prosthesis walkers than for the intact walker, especially during the stance phase of the intact leg (Figure 4). During stance of the intact leg, almost all gains were higher for the prosthesis walkers, but increases were more pronounced for the stance leg torques and least pronounced for the swing knee torque (Figure 4B). In particular, there was increased local and inter-joint feedback from the stance hip position in the prosthetic walkers. The pattern of how gains changed was similar for the different prosthesis walkers. Feedback gain trajectories can be found in the supplement (Figure S4). Feedback gains depended strongly on the sensory noise level and were five times higher at the lowest compared to the highest sensory noise level. The sensory noise level also had an effect on the relative magnitude of the gains. At low noise levels, walking with a prosthesis mainly resulted in an increase in feedback gains for the intact leg joints (and in particular the ankle) during stance (Figure S7A), while at the high noise level, walking with a prosthesis mainly resulted in an increase in feedback gains for the prosthesis knee and hip during stance (Figure S8A).

**Figure 4.**
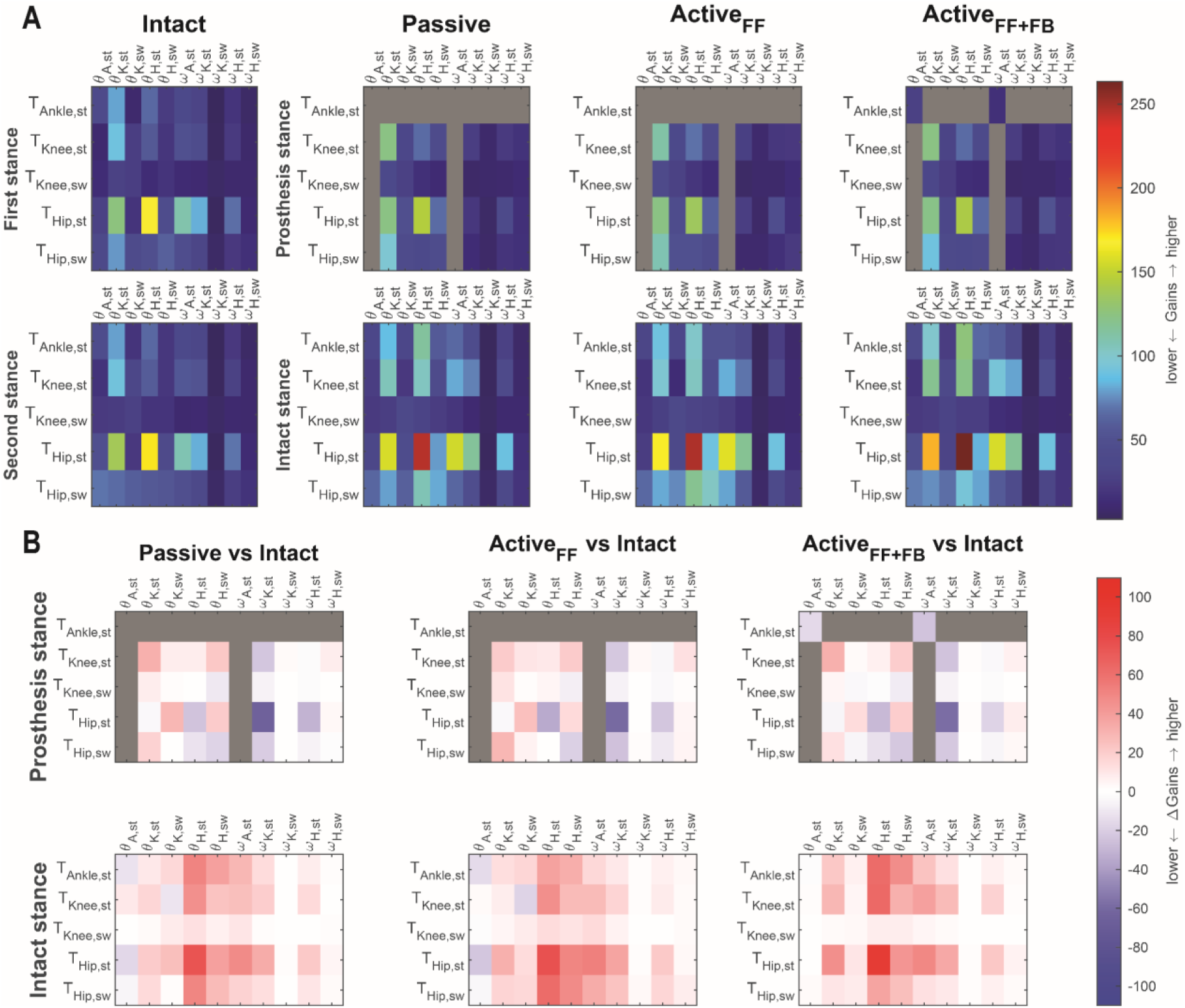
Root mean square feedback gains over a step (stance or swing) at a sensory noise level of 0.05^2^ (°)^2^s and 0.05^2^ (°/s)^2^s. Angular position gains are expressed in Nm/rad and angular velocity gains are expressed in Nms/rad. A. The first row presents feedback gains for the step in which the first leg of the intact walker is in stance (column 1) and the prosthesis leg for the prosthesis walkers is in stance (columns 2-4). The second row presents feedback gains for the step in which the second leg in the intact walker is in stance (column 1) and the intact leg in the prosthesis walkers is in stance (columns 2-4). Note that gains are identical for both steps for the intact walker. Gray blocks indicate feedback gains that are zero by design for the prosthesis models, i.e., ankle feedback gains (first row in each matrix) and ankle information driving the other joints (first and sixth columns in each matrix), with the exception of local ankle position and angular velocity gains for the feedforward and feedback-controlled prosthesis. B. Differences in feedback gains between the prosthesis walkers and intact walker for the first (prosthesis in stance, first row) and second step (intact leg in swing, second row). Positive values in red indicate larger gains for the prosthesis walkers, while blue values indicate larger gains for the intact walker. st refers to stance and sw to swing. For example, in the prosthetic stance plot, the hip_st_, knee_st_, and ankle_st_ refer to hip, knee, and ankle joints of the prosthesis leg in stance, while the hip_sw_ and knee_sw_ refer to the hip and knee joints of the intact leg in swing.

## Discussion

Our simulations suggest that walking with a passive prosthesis increases expected effort by 7% at low to 13% at high sensorimotor noise levels compared to walking with biological limbs. Increases in effort when walking with a passive prosthesis are due to both increases in feedforward effort, which are related to propulsion in the absence of noise, and expected feedback effort to stabilize walking against sensorimotor noise. A feedforward-controlled active ankle prosthesis restored expected effort in the prosthetic walker at low sensory noise levels by decreasing feedforward effort but was unable to reduce expected feedback torques, which might explain why expected effort remained higher than for the intact walker at high sensory noise levels. While the active ankle prosthesis with feedforward control and local ankle angle and angular velocity feedback reduced expected feedback torques compared to the other prosthesis walkers, the reduction in expected feedback effort was modest (i.e., some reduction at low but no reduction at high sensorimotor noise levels). This suggests that active ankle prostheses with feedforward and/or local feedback control may not restore the metabolic cost of walking when sensorimotor noise is considerable because they do not (sufficiently) restore the ankle’s contribution to stabilizing walking. We observed little differences in joint kinematic trajectories between the intact and prosthetic walkers, but the underlying control strategies were different. Feedback gains were larger for the prosthetic walkers, especially when the intact leg was in stance. Future work should investigate whether our overall observations are robust against modeling choices, i.e., whether they hold when using more detailed musculoskeletal models.

Our simulations predicted increased effort of walking with a passive prosthesis in line with experimental observations (1), but in contrast to previous simulation studies (26,28). Most simulation studies based on complex neuromusculoskeletal models predicted a similar metabolic cost for walking in individuals with a transtibial amputation walking with a passive prosthesis and in individuals without amputation (26,28), possibly because they did not account for the need to stabilize walking in the presence of sensorimotor noise. We accounted for sensorimotor noise but used a simple conceptual planar model. As a result, we could not directly assess metabolic energy but instead used the sum of expected torques squared as a measure of effort. Alternatively, positive mechanical work has been used as a measure of effort in simulations based on conceptual models (49). It is hard to tell which measure relates best to metabolic cost, mechanical work does not account for the metabolic energy consumed when muscles act isometrically and none of these measures accounts for differences in metabolic cost of eccentric and concentric muscle work. Nevertheless, the predicted increase in biological effort of 7%-13% (depending on the sensory noise level) when walking with a passive prosthesis instead of with biological limbs is in the order of magnitude of differences in the metabolic cost of walking that are observed between individuals with and without a transtibial amputation (1). Both higher feedforward and feedback effort contributed to the higher total effort in the passive prosthesis walker, which suggests that both effort for propulsion and stabilizing walking may contribute to the high metabolic cost of walking in individuals with a transtibial amputation. In our simulations, the differences in expected effort between the walker with a passive prosthesis and the intact walker were mostly due to differences in feedforward effort, but it is unlikely that we accurately captured the relative contribution of feedforward and feedback effort due to using a simple conceptual model. For example, we did not explicitly model the feet. Therefore, the passive prosthesis was modeled by a pin joint without any stiffness or damping, in contrast to conventional passive prostheses, which have well-tuned compliance that can contribute to the ankle torque through energy storage and return for propulsion and stabilization (50). Additionally, we did not model passive stiffness and damping in biological joints, whereas passive joint compliance contributes to stability (51). Further, we used a planar model, whereas stabilizing walking requires active control across planes (12,14). Our simulations thus suggest that both feedforward and feedback effort may contribute to the high metabolic cost of walking in individuals with a transtibial amputation using a passive prosthesis, but our approach is not suitable to quantitatively estimate the relative contribution of effort for propulsion versus stabilizing walking.

Our simulations suggest that active ankle prostheses can reduce the expected effort of walking by reducing the feedforward effort that biological joints have to generate but cannot fully eliminate compensation of the remaining biological joints for the loss of an ankle. The feedforward torques generate the mean trajectory in the absence of noise and are therefore responsible for forward propulsion of the body. Although feedforward or anticipatory strategies can be used to stabilize movement in the presence of uncertainty (22,52), our simulations predicted similar movement kinematics across sensory noise levels. Hence, we can interpret the reduction in biological feedforward effort as a reduced need for propulsion by the biological joints. Previous deterministic simulations (i.e., simulations that did not account for noise) also predicted that feedforward-controlled active prostheses can reduce the metabolic cost of walking compared to a passive prosthesis and even compared to individuals without a transtibial amputation (30). While active ankle prostheses have been observed to reduce the metabolic cost of walking compared to walking with a passive prosthesis (18,53), this is not generally the case (4,19,31). Our simulations likely overestimate the capacity of a feedforward-controlled active ankle prosthesis to reduce effort because they rely on the assumption that the human control policy is optimally adapted to walking with a prosthesis and that the prosthesis torque is optimally tuned to minimize effort. Here, we computed such an optimal prosthesis torque profile at a predefined constant walking speed. However, actuation of real-world devices often relies on feedback laws based on a limited number of prosthesis encoders and tuned to match data of walking mechanics, but is not optimally tuned to reduce effort (3,41). Handford and Srinivasan (29) compared optimized to real-world prosthesis actuation in simulation and indeed found that optimal torque profiles have the potential to reduce the metabolic cost of walking with respect to real-world prosthesis actuation.

Our simulations suggest that active ankle prostheses can reduce feedforward effort, but reductions in expected feedback effort were absent or modest, which may explain why active ankle prostheses may not restore the metabolic cost of walking. Active ankle prostheses reduced expected effort by reducing feedforward effort compared to the intact walker. Yet, the feedforward-controlled active prosthesis could not reduce the expected feedback effort. However, adding local feedback to the active feedforward-controlled prosthesis reduced the expected feedback effort, but only at the lowest sensorimotor noise levels. This suggests that the capacity of local feedback to reduce the effort related to stabilizing walking decreases as sensory information becomes less reliable. This is not surprising given that ankle torque corrections during walking in individuals without an amputation are poorly explained by local feedback (6). Instead, healthy individuals appear to modulate their ankle torque based on information about the whole body, such as center of mass position and velocity, suggesting that they integrate sensory information to stabilize walking (6). Hence, an active ankle prosthesis with full-state feedback might be required to reduce the effort for stabilizing walking when sensorimotor noise or environmental uncertainty is higher. Yet, such a full-state feedback-controlled active ankle prosthesis for stabilizing walking in individuals with an amputation may require many sensors and may therefore be infeasible for daily use. In addition, tuning the control parameters to different conditions, e.g., speeds or smoothness of the ground, might be challenging. In the future, simulations may be used to find control policies that minimize effort within the feasible set of sensory inputs (i.e., instrumentation) by exploring trade-offs between instrumentation and effort.

Our simulations further suggest that walking with a prosthesis has a limited effect on gait kinematics, in agreement with experimental observations (54,55), but has a large effect on the control policy. In line with our simulations, experimental studies found few differences in sagittal plane walking kinematics between individuals with and without a transtibial amputation (54,55). Yet, the few differences that are reported in literature, i.e., a lower range of motion in the prosthetic ankle (55) and more stance knee flexion in individuals with a transtibial amputation (54), were not present in our simulations. We might have artificially introduced similarity between the intact and prosthesis walkers by imposing step length and step time symmetry. As a result, our simulations could not adopt asymmetric step lengths or times, which have been observed in individuals with a transtibial amputation (56,57). Note that we also imposed a relatively slow walking speed of 0.6 m/s, as the average preferred walking speed of individuals with an amputation is between 0.8 and 1.0 m/s (1). Humans rely more on active control for stabilizing walking at slower than fast speeds (14), suggesting that walking at 0.6 m/s may impose greater stability demands than the walking speeds typically reported in the literature. Despite similar kinematics, the intact and prosthetic walkers achieved stable walking using different control strategies. Feedback gains were higher for the prosthetic walkers, especially when the intact leg was in stance. The increase in gain was especially large for local hip joint angle feedback of the intact leg during stance, suggesting that increased stiffness of the intact stance hip may be a compensation mechanism to stabilize walking after transtibial amputation. In contrast to our models, humans may not only achieve increased joint stiffness through sensorimotor feedback but also through agonist-antagonist co-activation. Individuals with a transtibial amputation were observed to walk with higher co-activation compared to individuals without an amputation for ankle and knee muscles, while co-activation has not been investigated in hip muscles (58). In addition, at the lowest sensory noise level, prosthesis walkers relied more on distal intact stance joints for feedback control compared to the intact walker, while at high sensory noise levels, the prosthesis walkers relied more on the proximal prosthesis leg joints over the stance phase (Figure S7 and Figure S8). Future studies are needed to test these model predictions (increased hip stiffness and a shift from distal to proximal compensation strategies with increasing noise) in individuals with a transtibial amputation.

Our simulations predict that joint kinematic variability is similar between individuals with and without a transtibial amputation at lower noise levels but larger in individuals with a transtibial amputation at higher noise levels, but experimental data to validate this result is lacking. Few papers report on intra-subject variability in joint kinematics in individuals with a transtibial amputation during unperturbed walking (59), and no studies compare joint kinematic variability between individuals with and without a transtibial amputation, which makes it challenging to validate our simulation results. Nevertheless, experimental observations suggest that swing foot clearance during walking is more variable in individuals with than in those without a transtibial amputation (60). The foot during swing can be considered the end-effector of the leg segments. Increased variability in joint angles may thus result in increased variability in foot position. Our simulations suggest that such increased variability in swing foot clearance might only be observed when uncertainty is high enough. Whether joint kinematic variability is indeed higher in individuals with compared to those without an amputation when uncertainty is high remains to be confirmed in future experimental studies.

In this work, we simulated walking by solving a deterministic approximation of a stochastic optimal control problem and not the exact stochastic problem. We approximated the state distribution by a Gaussian. In addition, we propagated the mean state using the nonlinear dynamics, while we propagated the covariance matrix based on a linear approximation of the dynamics, i.e., Lyapunov equations within a step and saltation matrix at impulsive contact. The resulting controllers stabilize the approximated dynamics but not necessarily the original dynamics. Linearizing the system dynamics to investigate the effect of noise on control of movement is common in the field of motor control (32,34,36). Such simulations have been shown to explain experimental observations, e.g., speed-accuracy trade-off in reaching movements (32), where including non-linear dynamics improved quantitative agreement with experimental data (35). Here, we did not linearize the dynamics of the mean state, which would not have been realistic for the highly nonlinear dynamics of walking, but we did linearize the dynamics of the state covariance around the mean state. Our results on the contribution of control for stabilizing walking to the energy cost of walking qualitatively matched experimental observations (1). Therefore, we think that simplified models that account for noise are useful to gain basic insights into how alterations in sensorimotor control affect the control of walking and thereby the metabolic cost. Yet, our approach is unsuitable for designing control policies that can be directly translated to real-world devices.

## Conclusion

Our simulations of walking in the presence of sensorimotor noise indicate that the cost of walking with a passive prosthesis is high due to both a lack of ankle feedforward propulsion and alterations in feedback control to stabilize walking. While an active ankle prosthesis may restore the cost of propulsion, it cannot restore the effort associated with stabilizing walking without sensory information beyond ankle kinematics. An active prosthesis with both feedforward and local feedback control reduces feedback-related effort but only modestly. The limited success of existing feedforward-controlled active prostheses in restoring the metabolic cost of walking in individuals with a transtibial amputation may thus be due to their limited ability to reduce the metabolic cost of stabilizing walking. Effectively reducing the metabolic cost in a broad group of prosthesis users might require active prostheses that support both propulsion and balance control.

## Competing interests

The authors declare no competing interests.

## Funding

The project was funded by a Fonds Wetenschappelijk Onderzoek (FWO) fellowship (1SF7322N) received by W.M.

## Contributions

The study was conceptualized and designed by WM, MA, RR, and FDG. Software implementation was handled by WM, TVW, and FDG. Simulations were conducted by WM. Data analysis and visualization were carried out by WM and FDG. WM, MA, RR, and FDG interpreted the results. The original draft was written by WM and FDG. WM, RR, MA, TVW, and FDG reviewed and edited the manuscript.

## Acknowledgements

We thank Lars D’Hondt and Dhruv Gupta for the insightful discussions about stochastic optimal control simulations of human movement.

## Notes

### Competing Interest Statement

The authors have declared no competing interest.

## References

1. Ettema S, Kal E, Houdijk H. General estimates of the energy cost of walking in people with different levels and causes of lower-limb amputation: a systematic review and meta-analysis. Prosthetics and Orthotics International. 14 oktober 2021;45(5):417–27. doi:10.1097/PXR.0000000000000035

2. Houdijk H, Pollmann E, Groenewold M, Wiggerts H, Polomski W. The energy cost for the step-to-step transition in amputee walking. Gait and Posture. 2009;30(1):35–40. doi:10.1016/j.gaitpost.2009.02.009 PubMed PMID: 19321343.

3. Montgomery JR, Grabowski AM. Use of a powered ankle–foot prosthesis reduces the metabolic cost of uphill walking and improves leg work symmetry in people with transtibial amputations. Journal of the Royal Society Interface. 2018;15(145). doi:10.1098/rsif.2018.0442 PubMed PMID: 30158189.

4. Quesada RE, Caputo JM, Collins SH. Increasing ankle push-off work with a powered prosthesis does not necessarily reduce metabolic rate for transtibial amputees. Journal of Biomechanics. 3 oktober 2016;49(14):3452-9. doi:10.1016/j.jbiomech.2016.09.015 PubMed PMID: 27702444.

5. Miller WC, Speechley M, Deathe B. The prevalence and risk factors of falling and fear of falling among lower extremity amputees. Archives of Physical Medicine and Rehabilitation. 1 augustus 2001;82(8):1031-7. doi:10.1053/apmr.2001.24295 PubMed PMID: 11494181.

6. Afschrift M, De Groote F, Jonkers I. Similar sensorimotor transformations control balance during standing and walking. PLOS Computational Biology. 25 juni 2021;17(6):e1008369. doi:10.1371/journal.pcbi.1008369

7. Hak L, Dieën JH van, Wurff P van der, Prins MR, Mert A, Beek PJ, e.a. Walking in an Unstable Environment: Strategies Used by Transtibial Amputees to Prevent Falling During Gait. Archives of Physical Medicine and Rehabilitation. 1 november 2013;94(11):2186–93. doi:10.1016/j.apmr.2013.07.020 PubMed PMID: 23916618.

8. Gates DH, Scott SJ, Wilken JM, Dingwell JB. Frontal plane dynamic margins of stability in individuals with and without transtibial amputation walking on a loose rock surface. Gait & Posture. 1 september 2013;38(4):570–5. doi:10.1016/j.gaitpost.2013.01.024

9. Cyr KM, Segal AD, Neptune RR, Klute GK. Biomechanical responses of individuals with transtibial amputation stepping on a coronally uneven and unpredictable surface. Journal of Biomechanics. 1 juni 2023;155:111622. doi:10.1016/j.jbiomech.2023.111622

10. Muijres W, Afschrift M, Ronsse R, De Groote F. Transtibial amputation increases the metabolic energy needed for stabilizing walking in the sagittal plane. Journal of Applied Physiology. juli 2026;141(1):71–84. doi:10.1152/japplphysiol.00877.2025

11. Segal AD, Klute GK. Lower-limb amputee recovery response to an imposed error in mediolateral foot placement. Journal of Biomechanics. 22 september 2014;47(12):2911–8. doi:10.1016/j.jbiomech.2014.07.008

12. Donelan JM, Shipman DW, Kram R, Kuo AD. Mechanical and metabolic requirements for active lateral stabilization in human walking. Journal of Biomechanics. 1 juni 2004;37(6):827-35. doi:10.1016/j.jbiomech.2003.06.002

13. O’Connor SM, Xu HZ, Kuo AD. Energetic cost of walking with increased step variability. Gait Posture. mei 2012;36(1):102–7. doi:10.1016/j.gaitpost.2012.01.014 PubMed PMID: 22459093; PubMed Central PMCID: PMC3372656.

14. Muijres W, Afschrift M, Ronsse R, De Groote F. Speeding up, not slowing down, decreases the metabolic energy needed to stabilize walking in the sagittal plane. J Appl Physiol. januari 2026;140(1):88–97. doi:10.1152/japplphysiol.00255.2025

15. Huang T wei P, Shorter KA, Adamczyk PG, Kuo AD. Mechanical and energetic consequences of reduced ankle plantar-flexion in human walking. Journal of Experimental Biology. 1 november 2015;218(22):3541–50. doi:10.1242/jeb.113910

16. Kuo AD. Energetics of actively powered locomotion using the simplest walking model. J Biomech Eng. februari 2002;124(1):113–20. doi:10.1115/1.1427703 PubMed PMID: 11871597.

17. Kuo AD, Donelan JM, Ruina A. Energetic Consequences of Walking Like an Inverted Pendulum: Step-to-Step Transitions. Exercise and Sport Sciences Reviews. april 2005;33(2):88.

18. Herr HM, Grabowski AM. Bionic ankle–foot prosthesis normalizes walking gait for persons with leg amputation. Proc Biol Sci. 7 februari 2012;279(1728):457-64. doi:10.1098/rspb.2011.1194 PubMed PMID: 21752817; PubMed Central PMCID: PMC3234569.

19. Gardinier ES, Kelly BM, Wensman J, Gates DH. A controlled clinical trial of a clinically-tuned powered ankle prosthesis in people with transtibial amputation. Clinical Rehabilitation. 2018;32(3):319–29. doi:10.1177/0269215517723054 PubMed PMID: 28750586.

20. Welker CG, Voloshina AS, Chiu VL, Collins SH. Shortcomings of human-in-the-loop optimization of an ankle-foot prosthesis emulator: a case series. Royal Society Open Science. 5 mei 2021;8(5):202020. doi:10.1098/rsos.202020

21. Ahuja S, Franz JR. The metabolic cost of walking balance control and adaptation in young adults. Gait & Posture. 1 juli 2022;96:190-4. doi:10.1016/j.gaitpost.2022.05.031

22. Voloshina AS, Kuo AD, Daley MA, Ferris DP. Biomechanics and energetics of walking on uneven terrain. Journal of Experimental Biology. november 2013;216(21):3963–70. doi:10.1242/jeb.081711 PubMed PMID: 23913951.

23. Houdijk H, Blokland IJ, Nazier SA, Castenmiller SV, van den Heuvel I, IJmker T. Effects of Handrail and Cane Support on Energy Cost of Walking in People With Different Levels and Causes of Lower Limb Amputation. Archives of Physical Medicine and Rehabilitation. 1 juli 2021;102(7):1340-1346.e3. doi:10.1016/j.apmr.2021.02.007

24. IJmker T, Noten S, Lamoth CJ, Beek PJ, van der Woude LHV, Houdijk H. Can external lateral stabilization reduce the energy cost of walking in persons with a lower limb amputation? Gait & Posture. 1 september 2014;40(4):616–21. doi:10.1016/j.gaitpost.2014.07.013

25. Koelewijn AD, van den Bogert AJ. Joint contact forces can be reduced by improving joint moment symmetry in below-knee amputee gait simulations. Gait & Posture. 1 september 2016;49:219–25. doi:10.1016/j.gaitpost.2016.07.007

26. Miller RH, Russell Esposito E. Transtibial limb loss does not increase metabolic cost in three-dimensional computer simulations of human walking. PeerJ. 4 augustus 2021;9:e11960. doi:10.7717/PEERJ.11960

27. Fey NP, Klute GK, Neptune RR. Optimization of Prosthetic Foot Stiffness to Reduce Metabolic Cost and Intact Knee Loading During Below-Knee Amputee Walking: A Theoretical Study. J Biomech Eng. 26 oktober 2012;134(111005). doi:10.1115/1.4007824

28. Falisse A, Serrancolí G, Dembia CL, Gillis J, Jonkers I, De Groote F. Rapid predictive simulations with complex musculoskeletal models suggest that diverse healthy and pathological human gaits can emerge from similar control strategies. Journal of the Royal Society Interface. 1 augustus 2019;16(157). doi:10.1098/rsif.2019.0402 PubMed PMID: 31431186.

29. Handford ML, Srinivasan M. Energy-optimal human walking with feedback-controlled robotic prostheses: A computational study. IEEE Transactions on Neural Systems and Rehabilitation Engineering. 1 september 2018;26(9):1773–82. doi:10.1109/TNSRE.2018.2858204 PubMed PMID: 30040647.

30. Handford ML, Srinivasan M. Robotic lower limb prosthesis design through simultaneous computer optimizations of human and prosthesis costs. Scientific Reports. 9 februari 2016;6(1):1-7. doi:10.1038/srep19983 PubMed PMID: 26857747.

31. Kim J, Wensman J, Colabianchi N, Gates DH. The influence of powered prostheses on user perspectives, metabolics, and activity: a randomized crossover trial. Journal of NeuroEngineering and Rehabilitation. 16 maart 2021;18(1):49. doi:10.1186/s12984-021-00842-2

32. Harris CM, Wolpert DM. Signal-dependent noise determines motor planning. Nature. 20 augustus 1998;394(6695):780-4. doi:10.1038/29528 PubMed PMID: 9723616.

33. Todorov E, Jordan MI. Optimal feedback control as a theory of motor coordination. Nature Neuroscience. 2002;5(11):1226–35. doi:10.1038/nn963 PubMed PMID: 12404008.

34. Diedrichsen J. Optimal Task-Dependent Changes of Bimanual Feedback Control and Adaptation. Current Biology. 9 oktober 2007;17(19):1675-9. doi:10.1016/j.cub.2007.08.051 PubMed PMID: 17900901.

35. Van Wouwe T, Ting LH, De Groote F. An approximate stochastic optimal control framework to simulate nonlinear neuro-musculoskeletal models in the presence of noise. PLoS computational biology. 1 juni 2022;18(6):e1009338. doi:10.1371/journal.pcbi.1009338 PubMed PMID: 35675227.

36. Kuo AD. An optimal state estimation model of sensory integration in human postural balance [Internet]. 1 september 2005 [geciteerd 12 juli 2023]. Beschikbaar op: http://deepblue.lib.umich.edu/handle/2027.42/49185

37. Koelewijn AD, van den Bogert AJ. A solution method for predictive simulations in a stochastic environment. Journal of Biomechanics. 7 mei 2020;104:109759. doi:10.1016/j.jbiomech.2020.109759

38. Kelly M. An introduction to trajectory optimization: How to do your own direct collocation∗. SIAM Review. 2017;59(4):849–904. doi:10.1137/16M1062569

39. Anderson FC, Pandy MG. A Dynamic Optimization Solution for Vertical Jumping in Three Dimensions. Computer Methods in Biomechanics and Biomedical Engineering. 1 januari 1999;2(3):201-31. doi:10.1080/10255849908907988 PubMed PMID: 11264828.

40. Delp SL, Loan JP, Hoy MG, Zajac FE, Topp EL, Rosen JM. An interactive graphics-based model of the lower extremity to study orthopaedic surgical procedures. IEEE Trans Biomed Eng. augustus 1990;37(8):757–67. doi:10.1109/10.102791 PubMed PMID: 2210784.

41. Eilenberg MF, Geyer H, Herr H. Control of a powered ankle-foot prosthesis based on a neuromuscular model. IEEE Transactions on Neural Systems and Rehabilitation Engineering. april 2010;18(2):164–73. doi:10.1109/TNSRE.2009.2039620 PubMed PMID: 20071268.

42. Markowitz J, Krishnaswamy P, Eilenberg MF, Endo K, Chris CB, Herr H. Speed adaptation in a powered transtibial prosthesis controlled with a neuromuscular model. Philosophical Transactions of the Royal Society B: Biological Sciences. 2011;366(1570):1621–31. doi:10.1098/rstb.2010.0347

43. Shi F, Rymer WZ, Son J. Ankle Joint Angle Influences Relative Torque Fluctuation during Isometric Plantar Flexion. Bioengineering (Basel). 18 maart 2023;10(3):373. doi:10.3390/bioengineering10030373 PubMed PMID: 36978764; PubMed Central PMCID: PMC10045061.

44. Mazara N, Hess AJ, Chen J, Power GA. Activation reduction following an eccentric contraction impairs torque steadiness in the isometric steady-state. J Sport Health Sci. juli 2018;7(3):310–7. doi:10.1016/j.jshs.2018.05.001 PubMed PMID: 30356642; PubMed Central PMCID: PMC6189235.

45. Wolpert DM, Miall RC, Kawato M. Internal models in the cerebellum. Trends in Cognitive Sciences. 1 september 1998;2(9):338–47. doi:10.1016/S1364-6613(98)01221-2 PubMed PMID: 21227230.

46. Kong NJ, Joe Payne J, Zhu J, Johnson AM. Saltation Matrices: The Essential Tool for Linearizing Hybrid Dynamical Systems. Proceedings of the IEEE. juni 2024;112(6):585–608. doi:10.1109/JPROC.2024.3440211

47. Andersson JAE, Gillis J, Horn G, Rawlings JB, Diehl M. CasADi: a software framework for nonlinear optimization and optimal control. Mathematical Programming Computation. 14 maart 2019;11(1):1-36. doi:10.1007/s12532-018-0139-4

48. Wächter A, Biegler LT. On the implementation of an interior-point filter line-search algorithm for large-scale nonlinear programming. Mathematical Programming. 2006;106(1):25–57. doi:10.1007/s10107-004-0559-y

49. Srinivasan M, Ruina A. Computer optimization of a minimal biped model discovers walking and running. Nature. januari 2006;439(7072):72–5. doi:10.1038/nature04113

50. Zelik KE, Collins SH, Adamczyk PG, Segal AD, Klute GK, Morgenroth DC, e.a. Systematic Variation of Prosthetic Foot Spring Affects Center-of-Mass Mechanics and Metabolic Cost During Walking. IEEE Transactions on Neural Systems and Rehabilitation Engineering. augustus 2011;19(4):411–9. doi:10.1109/TNSRE.2011.2159018

51. John CT, Anderson FC, Higginson JS, Delp SL. Stabilisation of walking by intrinsic muscle properties revealed in a three-dimensional muscle-driven simulation. Computer Methods in Biomechanics and Biomedical Engineering. april 2013;16(4):451–62. doi:10.1080/10255842.2011.627560 PubMed PMID: 22224406.

52. Gates DH, Wilken JM, Scott SJ, Sinitski EH, Dingwell JB. Kinematic strategies for walking across a destabilizing rock surface. Gait Posture. januari 2012;35(1):36–42. doi:10.1016/j.gaitpost.2011.08.001 PubMed PMID: 21890361; PubMed Central PMCID: PMC3262902.

53. Russell Esposito E, Aldridge Whitehead JM, Wilken JM. Step-to-step transition work during level and inclined walking using passive and powered ankle-foot prostheses. Prosthet Orthot Int. juni 2016;40(3):311–9. doi:10.1177/0309364614564021 PubMed PMID: 25628378.

54. Kovac I, Medved V, Kasović M, Ž H, Luzar-Stiffler V, Pecina M. Instrumented joint mobility analysis in traumatic transtibial amputee patients. Periodicum Biologorum. 1 maart 2010;112:25-31.

55. Gates DH, Dingwell JB, Scott SJ, Sinitski EH, Wilken JM. Gait characteristics of individuals with transtibial amputations walking on a destabilizing rock surface. Gait and Posture. mei 2012;36(1):33–9. doi:10.1016/j.gaitpost.2011.12.019 PubMed PMID: 22469772.

56. Hak L, Van Dieën JH, Van Der Wurff P, Houdijk H. Stepping asymmetry among individuals with unilateral transtibial limb loss might be functional in terms of gait stability. Physical Therapy. 1 oktober 2014;94(10):1480-8. doi:10.2522/ptj.20130431 PubMed PMID: 24903115.

57. Wedge RD, Sup FC, Umberger BR. Metabolic cost of transport and stance time asymmetry in individuals with unilateral transtibial amputation using a passive prostheses while walking. Clinical Biomechanics. 1 april 2022;94:105632. doi:10.1016/j.clinbiomech.2022.105632

58. Seyedali M, Czerniecki JM, Morgenroth DC, Hahn ME. Co-contraction patterns of trans-tibial amputee ankle and knee musculature during gait. J Neuroeng Rehabil. 28 mei 2012;9:29. doi:10.1186/1743-0003-9-29 PubMed PMID: 22640660; PubMed Central PMCID: PMC3480942.

59. Gates DH, Darter BJ, Dingwell JB, Wilken JM. Comparison of walking overground and in a Computer Assisted Rehabilitation Environment (CAREN) in individuals with and without transtibial amputation. J Neuroeng Rehabil. 14 november 2012;9:81. doi:10.1186/1743-0003-9-81 PubMed PMID: 23150903; PubMed Central PMCID: PMC3543217.

60. De Asha RA. Variability and Distribution of Minimum Toe Clearance in Individuals with Unilateral Transtibial Amputation: The Effects of Walking Speed. JPO: Journal of Prosthetics and Orthotics. juli 2015;27(3):78. doi:10.1097/JPO.0000000000000062

