## Supplementary text for "Simulations predict that alterations in balance control contribute to the higher cost of walking with a prosthesis"

### Supplementary material

#### S1. Finding a Sensory noise level capturing experimental data

We compared outcomes of OCP solutions of the intact walker with different sensory noise levels with experimental data of unperturbed walking of healthy individuals (1) to find a representative sensory noise level. We compared the between-stride variability in fore-aft center of mass (CoM) position and ankle torques between OCP solutions and experimental data. We hypothesized that these outcomes would be sensitive to the sensory noise levels, as our previous findings suggested that humans modulate ankle torques based on CoM kinematics to stabilize walking against perturbations (2).

We compared results from simulations against CoM kinematic and ankle torque data from 18 healthy young adults who walked on a treadmill while marker trajectories and ground reaction force data were collected (1). Joint kinematics were calculated using OpenSim's Inverse Kinematics Tool based on a model that was scaled to the subjects' anthropometry. Then, CoM kinematics and ankle torques were calculated using the OpenSim BodyKinematics analysis and the Inverse Dynamics Tool. Time series were normalized to a stride based on left heel strike events and torques were normalized to body mass. Additional information on the processing of experimental data (i.e., marker trajectories and ground reaction forces) can be found in (1). We computed CoM variability for the OCP solutions based on a linear approximation. In other words, we computed the variability in CoM kinematics as the product of the partial derivative of the CoM kinematics with respect to the states and the state covariance. We compared torque variability from experiments with feedback torques obtained from the optimal control problem, after normalization to body mass. Since intact walking is approximately symmetric, we compared simulation and experimental results over a step.

Our analysis did not provide a clear indication as to which sensory noise level best represented experimental data (Figure S1). The pattern of ankle torque variability as a function of the step differed between simulations and experiments, probably because we did not model the double support phase. Simulated mean ankle torque variability in simulation was not very sensitive to noise level and was smaller – yet had the same order of magnitude - in simulation (0.06 Nm/kg) than experimentally observed (0.10 Nm/kg). Therefore, this criterion did not clearly discriminate between the realism of sensory noise levels. Also, the pattern of CoM position variability as a function of the step differed between simulations and experiments. While simulated and experimental CoM position variability were similar early in the step, the simulated CoM position variability increased towards the end of the step, but the experimental CoM position variability did not. This increase in variability during the step was larger for the highest sensory noise level tested here. However, this criterion did not clearly discriminate between the realism of the other sensory noise levels. Given the limited effect of sensory noise level on the agreement in ankle torque and CoM position variability trajectories between simulations and experimental data, we decided to report the solutions for the median sensory noise level (of 0.05 $(°)^2\mathrm{s}$ and 0.05 $\left(\frac{°}{\mathrm{s}}\right)^2 \mathrm{s}$) in the main manuscript.

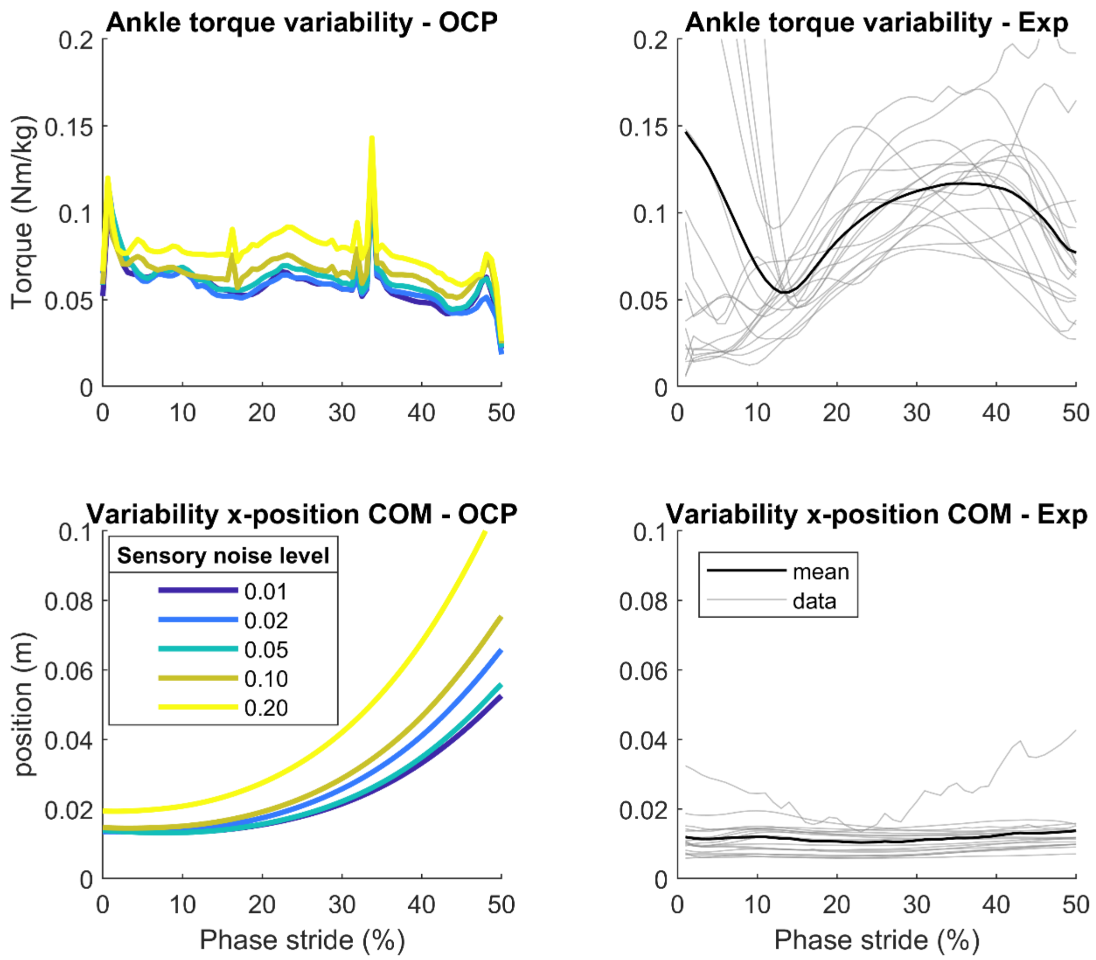


*Figure S1. Variability in ankle torques (top row) and fore-aft CoM position (bottom row) in OCP solutions (first column) and experiments (second column). Simulation results for the intact walker walking with different levels of sensory noise are shown left, while experimental data from 18 healthy young adults walking on a treadmill without perturbation is shown on the right (1). Experimental data were normalized to left heel-strike events and presented for the first 50% of the stride (i.e., including double support).*

#### S2. Joint angular velocities

We observed few differences in the mean joint angular velocities between the intact and prosthesis walkers (Figure S2). Compared to the intact walker, the variability in intact knee joint angular velocities over the first half of the swing phase was larger in the passive prosthesis and the feedforward control active prosthesis walkers. Yet, variability in joint angular velocities was similar across other joints and gait phases. In addition, we observed that variability in joint angular velocities was similar between the feedforward and local feedback-controlled active prosthesis walker and the intact walker.

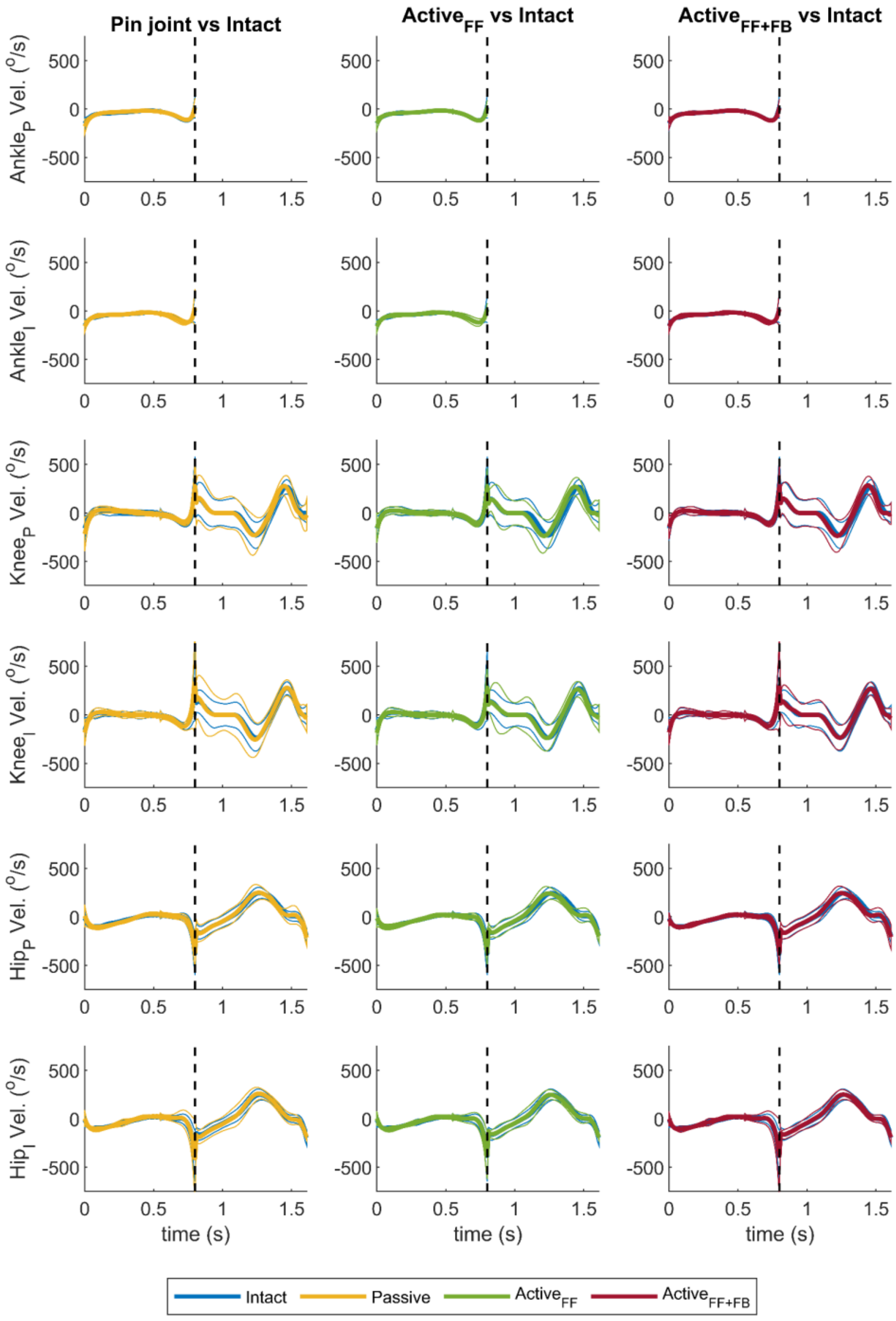

*Figure S2. Joint angular velocities for prosthesis walkers versus the intact walker at a sensory noise level of 0.05 $(°)^2$s and 0.05 $(°/s)^2$s. The optimal control solutions of the passive prosthesis are shown in yellow, of the feedforward-controlled active prosthesis in green, the feedforward and local feedback-controlled active prosthesis in red, and the intact walker in blue. Joint angular velocities are displayed for the ankles, knees, and hips of the intact (I) and prosthetic (P) leg. Thick and thin lines represent the mean and the mean +/- 1 standard deviation, respectively. The dashed vertical line indicates the moment of foot contact, separating the stance phase from the swing phase. Note that the transition between these phases is discontinuous due to impulsive contact.*

#### S3. Joint torque and feedback gain trajectories

##### *Joint torques*

Joint torques – mean and variability – were similar for the intact walkers and prosthesis walkers (Figure S3). The ankle torques of the prosthesis leg were different from those of the intact walker due to differences in actuation and control; see the description of the control policy for the prosthesis walkers in the Methods section. Additionally, we observed slightly higher variability in swing phase hip torques in the prosthesis than in the intact walkers. Overall, mean torques and torque variability were very similar between the prosthesis and intact walkers for the biological joints across the stance and swing phase.

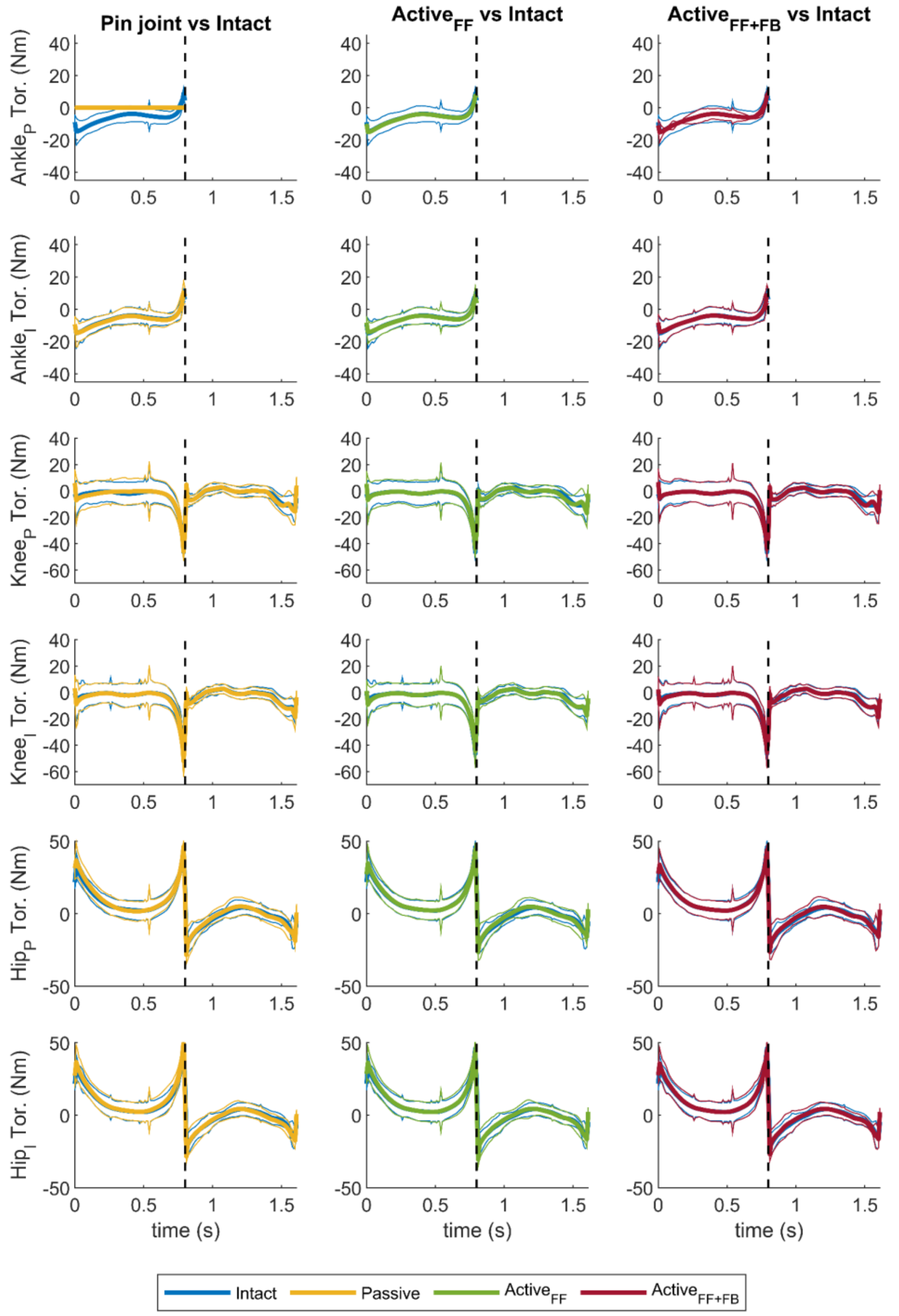


*Figure S3. Feedforward and expected feedback torques for prosthesis walkers versus the intact walker at a sensory noise level of 0.05 $(°)^2$s and 0.05 $(°/s)^2$s. The optimal control solutions of the passive prosthesis are shown in yellow, of the feedforward controlled active prosthesis in green, the feedforward and feedback controlled active prosthesis in red, and the intact walker in blue. Joint torques are displayed for the ankles, knees, and hips of the intact (I) and prosthetic (P) leg. Thick and thin lines represent the mean and the mean +/- 1 standard deviation, respectively. The dashed vertical line indicates the moment of toe-off, separating the stance phase from the swing phase. Note that feedforward torques are represented by the mean torque, while feedback torques are represented by the standard deviation around the mean. In contrast to the prosthetic ankle of the passive prosthesis walker that does not generate feedforward or feedback torques, the active prosthesis ankle generates feedforward torques in the feedforward-controlled active prosthesis walker and feedback and feedforward torques for the feedforward-controlled active prosthesis walker with local feedback.*

##### *Feedback gains*

The magnitude of feedback gains was larger in the prosthesis walkers than in the intact walker over the intact leg's stance phase (Figure S4). The magnitude of inter-joint feedback gains driving stance leg joints was larger in the prosthesis walker than in the intact walker in the intact limb (Figure S4B), and was highest for hip joint position information. In addition, feedback gain magnitudes were lower over the step in which the prosthesis leg was in stance (Figure S4A) versus the step in which the intact leg was in stance (Figure S4B).

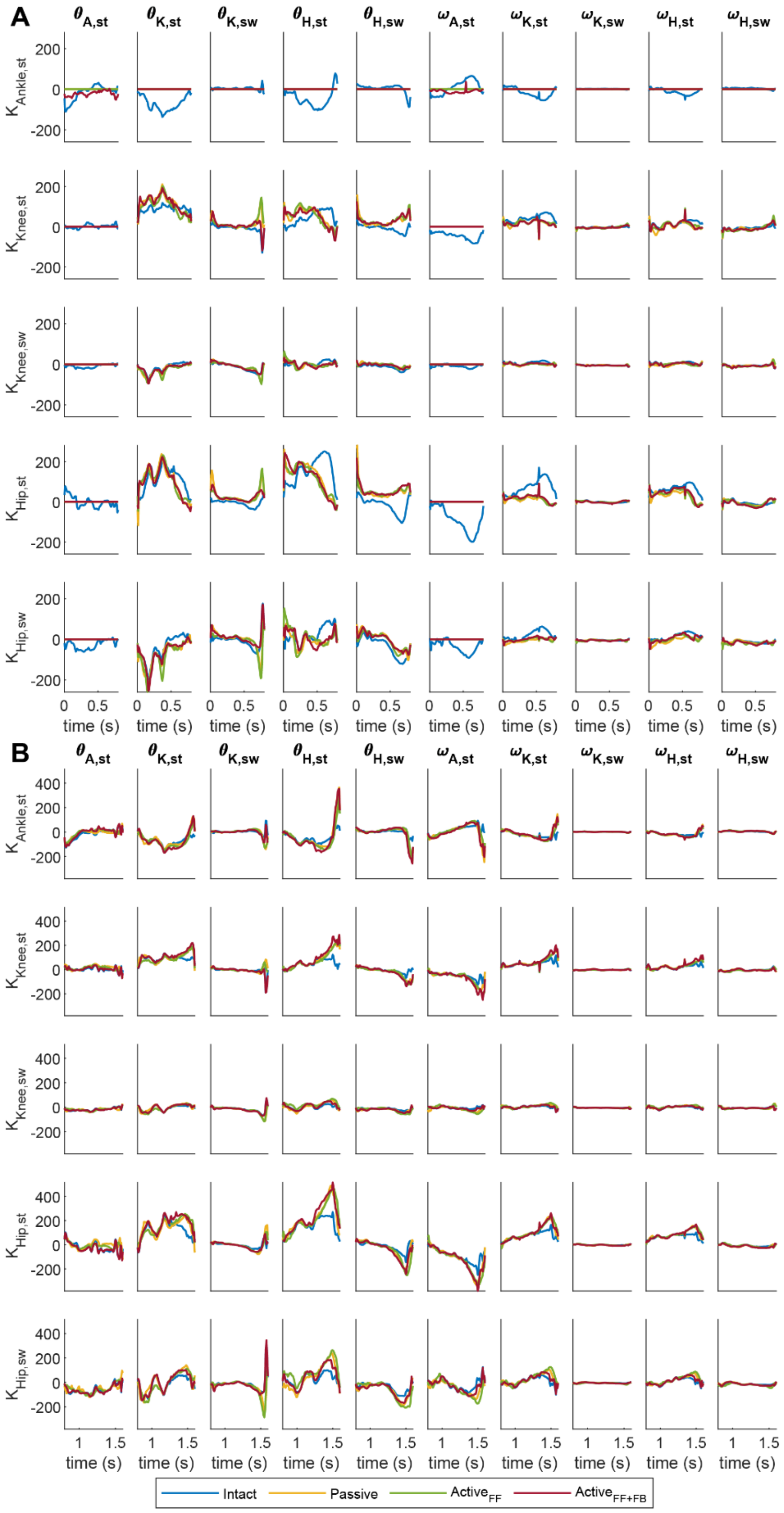


*Figure S4. Feedback gain trajectories for the intact and prosthesis walkers at a sensory noise level of 0.05 $(^o)^2$s and 0.05 $(^o/s)^2$s. Joints (five in total) received time-varying linear feedback from joint angles and angular velocities (in columns). A. Feedback gain trajectories for the first step, in which, for prosthesis walkers, the prosthesis leg is in stance. We modeled the effect of walking with a prosthesis on feedback control by setting ankle feedback gains (first row) and ankle information driving the other joints (first and sixth columns) to zero. In the case of the feedforward-controlled prosthesis with local feedback, ankle position and angular velocity gains could be non-zero. B. Feedback gain trajectories for the second step, in which, for the prosthesis walkers, the intact leg was in stance.*

#### S4. Effect of sensory noise

##### *Joint angles*

With increasing sensory noise levels, differences in mean swing leg kinematics between the intact walker and prosthetic walkers increased (Figure S5 and Figure S6). Differences between intact and prosthesis walkers were primarily driven by greater adaptations in the intact walkers. Specifically, intact walkers showed larger increases in swing-phase knee flexion in both legs and a greater increase in terminal-swing hip flexion than the prosthetic leg of prosthesis walkers. In addition, variability in the intact stance ankle angle increased with higher sensory noise and was larger in the prosthesis walkers in terminal stance in the intact leg.

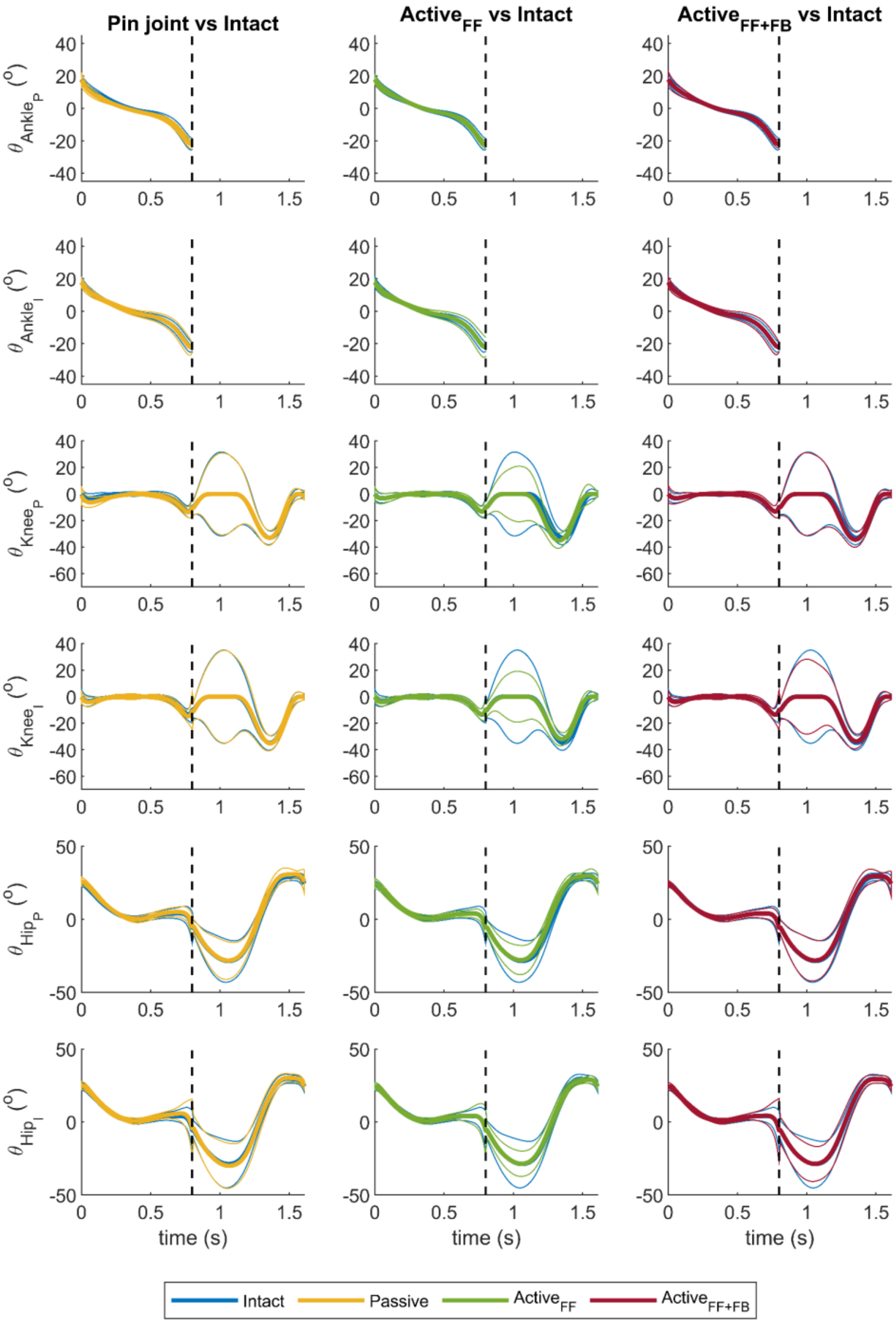


*Figure S5. Joint angular positions ($\theta$) over a stride for the prosthesis walkers versus the intact walker at a sensory noise level of $0.01^2$ $(°)^2$s and $0.01^2$ $(°/s)^2$s. The optimal control solutions of the passive prosthesis are shown in yellow, of the feedforward controlled active prosthesis in green, the feedforward and feedback controlled active prosthesis in red, and the intact walker in blue for the intact (I) and the prosthetic (P) legs. Joint angular positions are displayed for the ankles, knees, and hips of the intact and prosthetic legs. Thick and thin lines represent the mean and the mean +/- 1 standard deviation, respectively. The dashed vertical line indicates the moment of toe-off, separating the stance phase from the swing phase.*

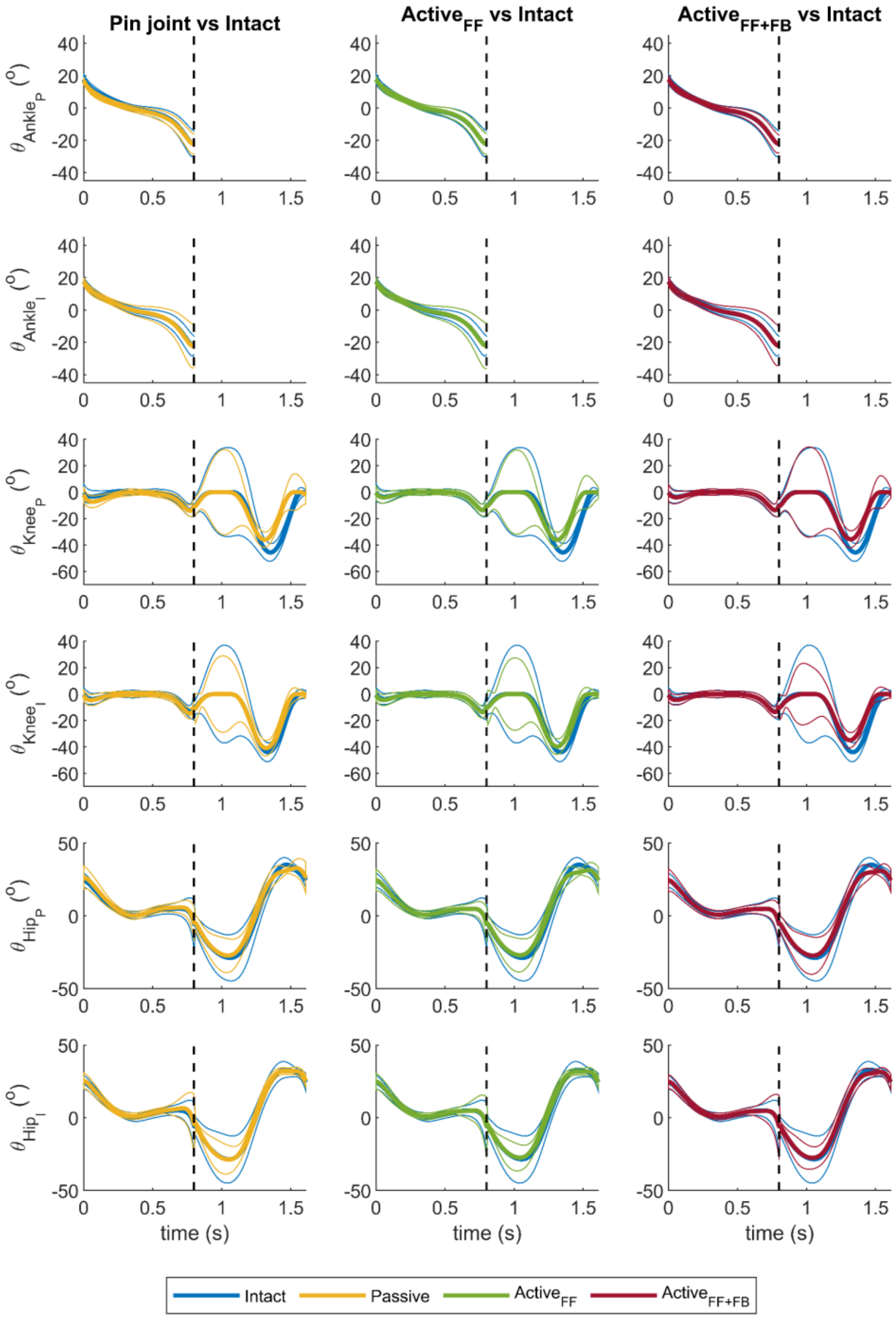


*Figure S6. Joint angular positions ($\theta$) over a stride for the prosthesis walkers versus the intact walker at a sensory noise level of $0.2^2$ $(°)^2$s and $0.2^2$ $(°/s)^2$s. The optimal control solutions of the passive prosthesis are shown in yellow, of the feedforward controlled active prosthesis in green, the feedforward and feedback controlled active prosthesis in red, and the intact walker in blue for the intact (I) and the prosthetic (P) legs. Joint angular positions are displayed for the ankles, knees, and hips of the intact and prosthetic legs. Thick and thin lines represent the mean and the mean +/- 1 standard deviation, respectively. The dashed vertical line indicates the moment of toe-off, separating the stance phase from the swing phase.*

##### *Feedback gains*

Overall, feedback gains were higher for the prosthesis walker than for the intact walker, but how much and which feedback gains were higher in the prosthetic walkers depended on the sensory noise levels (Figure S7 and Figure S8). At the lowest noise level, position feedback gains from joints on the intact leg over the stance phase were higher in the prosthesis walkers than in the intact walker (Figure S7). In particular, local and inter-joint feedback gains were higher across joints of the intact leg over the stance phase for prosthesis walkers (Figure S7B). At high noise levels, we observed higher feedback gains driving the proximal joints of the prosthesis leg (knee and hip) in stance for prosthesis walkers than for intact walker (Figure S8B).

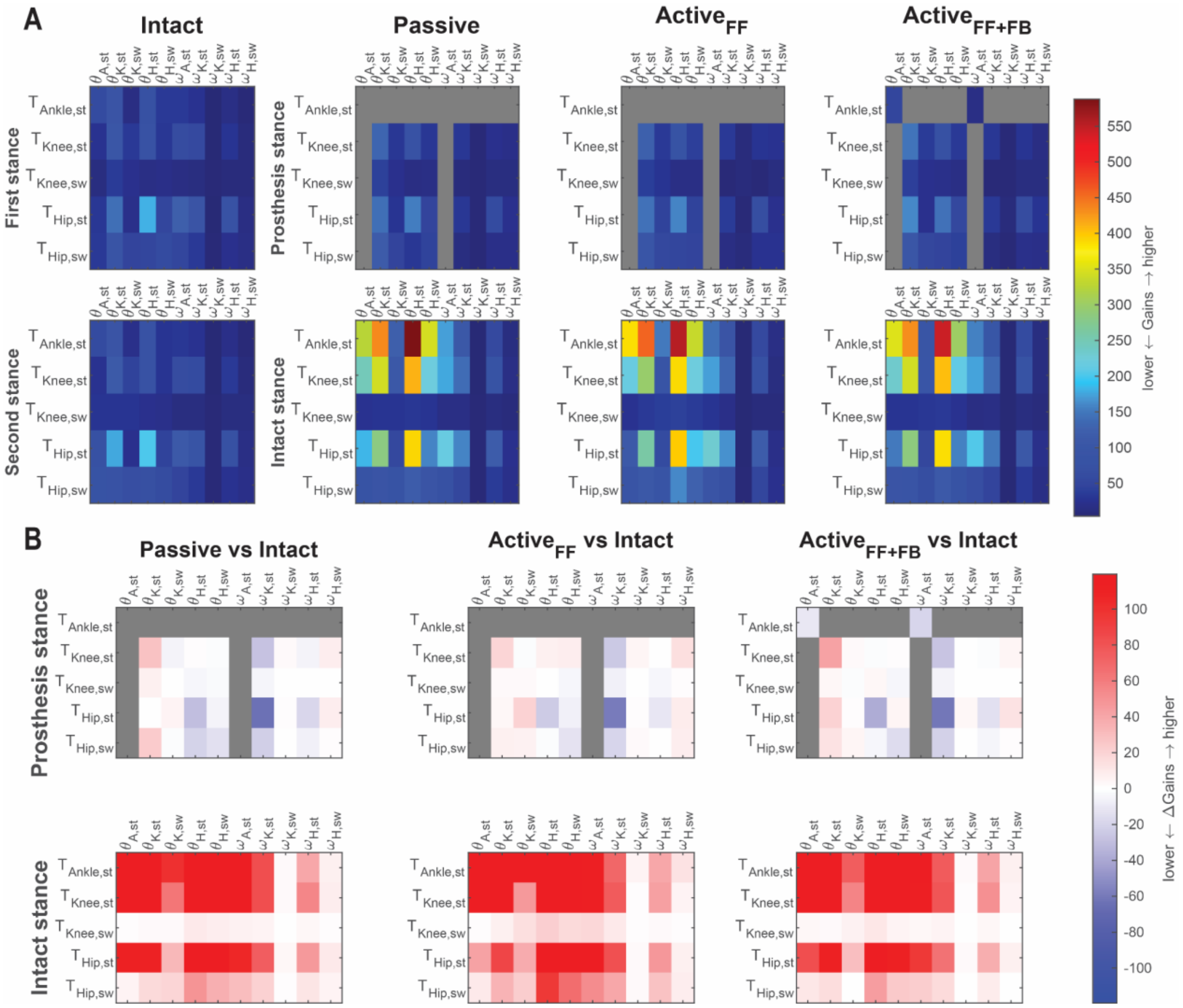


*Figure S7. Root mean square feedback gains over a step (stance or swing) at a sensory noise level of 0.01$^2$ (°)$^2$s and 0.01$^2$ (°/s)$^2$s. Angular position gains are expressed in Nm/rad and angular velocity gains are expressed in Nms/rad.* ***A.*** *The first row presents feedback gains for the step in which the first leg of the intact walker is in stance (column 1) and the prosthesis leg for the prosthesis walkers is in stance (columns 2-4). The second row presents feedback gains for the step in which the second leg in the intact walker is in stance (column 1) and the intact leg in the prosthesis walkers is in stance (columns 2-4). Note that gains are identical for both steps for the intact walker. Gray blocks indicate feedback gains that are zero by design for the prosthesis models, i.e., ankle feedback gains (first row in each matrix) and ankle information driving the other joints (first and sixth columns in each matrix), with the exception of local ankle position and angular velocity gains for the feedforward and feedback-controlled prosthesis.* ***B.*** *Differences in feedback gains between the prosthesis walkers and intact walker for the first (versus the prosthesis stance leg, first row) and second step (versus the intact stance leg, second row). Note that we used y-labels of the prosthesis walkers from A to indicate the comparison between legs from the intact versus the prosthesis walkers. Positive values in red indicate larger gains for the prosthesis walkers, while blue values indicate larger gains for the intact walker. st refers to stance and sw to swing. For example, in the prosthesis stance plots the hip$_{st}$, knee$_{st}$, and ankle$_{st}$ refer to hip, knee, and ankle joints of the prosthesis leg in stance, while the hip$_{sw}$ and knee$_{sw}$ refer to the hip and knee joints of the intact leg in swing.*

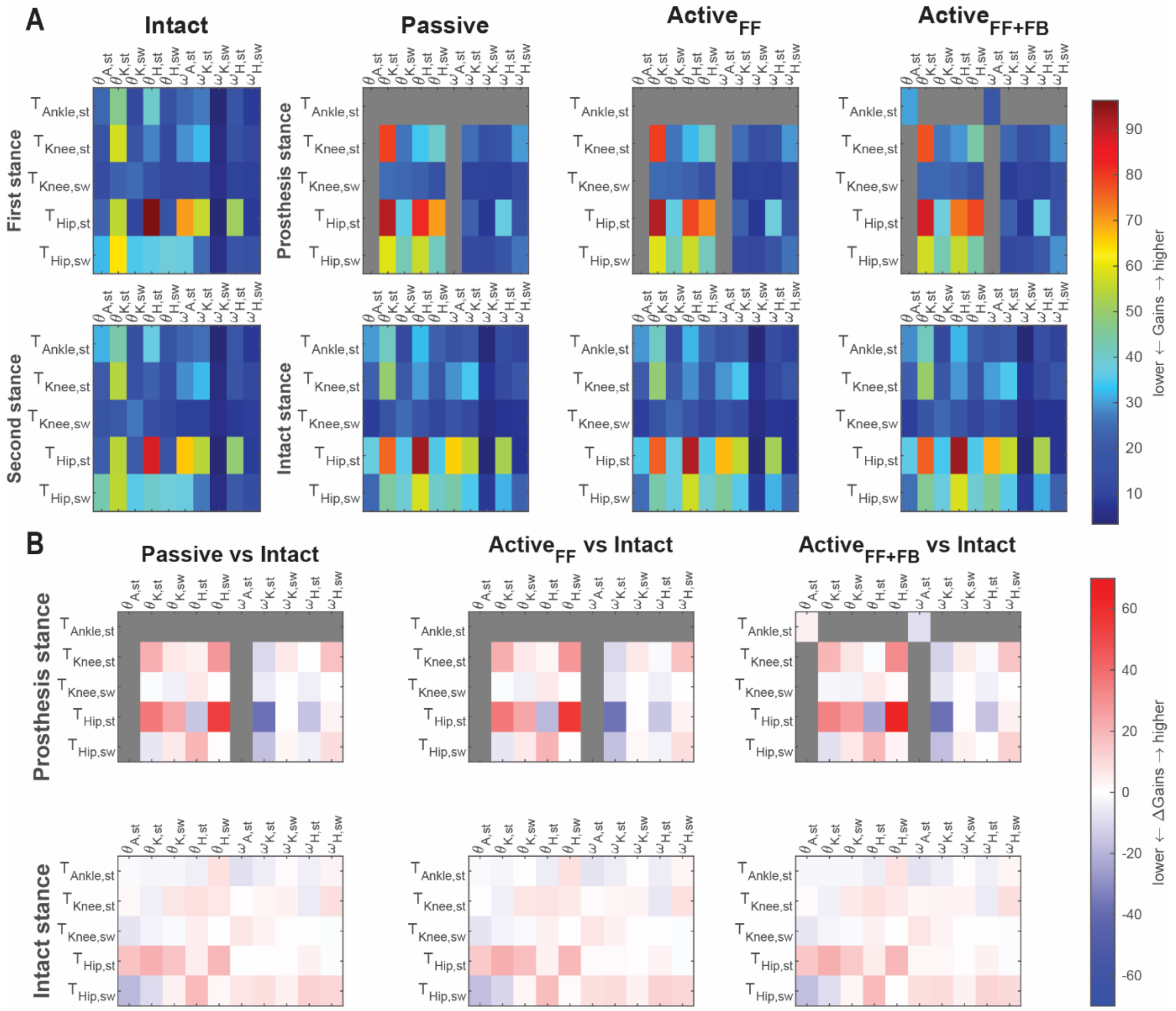


*Figure S8. Root mean square feedback gains over a step (stance or swing) at a sensory noise level of $0.2^2$ $(°)^2$s and $0.2^2$ $(°/s)^2$s. Angular position gains are expressed in Nm/rad and angular velocity gains are expressed in Nms/rad.* ***A.*** *The first row presents feedback gains for the step in which the first leg of the intact walker is in stance (column 1) and the prosthesis leg for the prosthesis walkers is in stance (columns 2-4). The second row presents feedback gains for the step in which the second leg in the intact walker is in stance (column 1) and the intact leg in the prosthesis walkers is in stance (columns 2-4). Note that gains are identical for both steps for the intact walker. Gray blocks indicate feedback gains that are zero by design for the prosthesis models, i.e., ankle feedback gains (first row in each matrix) and ankle information driving the other joints (first and sixth columns in each matrix), with the exception of local ankle position and angular velocity gains for the feedforward and feedback-controlled prosthesis.* ***B.*** *Differences in feedback gains between the prosthesis walkers and intact walker for the first (versus the prosthesis stance leg, first row) and second step (versus the intact stance leg, second row). Note that we used y-labels of the prosthesis walkers from A to indicate the comparison between legs from the intact versus the prosthesis walkers. Positive values in red indicate larger gains for the prosthesis walkers, while blue values indicate larger gains for the intact walker. st refers to stance and sw to swing. For example, in the prosthesis stance plots the $hip_{st}$, $knee_{st}$, and $ankle_{st}$ refer to hip, knee, and ankle joints of the prosthesis leg in stance, while the $hip_{sw}$ and $knee_{sw}$ refer to the hip and knee joints of the intact leg in swing.*

### References


1. Afschrift M, van Deursen R, De Groote F, Jonkers I. Increased use of stepping strategy in response to medio-lateral perturbations in the elderly relates to altered reactive tibialis anterior activity. Gait & Posture. 2019 Feb 1;68:575–82. doi:10.1016/j.gaitpost.2019.01.010

2. Afschrift M, De Groote F, Jonkers I. Similar sensorimotor transformations control balance during standing and walking. PLOS Computational Biology. 2021 Jun 25;17(6):e1008369. doi:10.1371/journal.pcbi.1008369
